# Chromatin phase separation and nuclear shape fluctuations are correlated in a polymer model of the nucleus

**DOI:** 10.1101/2023.12.16.571697

**Authors:** Ali Goktug Attar, Jaroslaw Paturej, Edward J. Banigan, Aykut Erbas

## Abstract

Abnormalities in the shapes of mammalian cell nuclei are hallmarks of a variety of diseases, including progeria, muscular dystrophy, and various cancers. Experiments have shown that there is a causal relationship between chromatin organization and nuclear morphology. Decreases in heterochromatin levels, perturbations to heterochromatin organization, and increases in euchromatin levels all lead to misshapen nuclei, which exhibit deformations, such as nuclear blebs and nuclear ruptures. However, the polymer physical mechanisms of how chromatin governs nuclear shape and integrity are poorly understood. To investigate how heterochromatin and euchromatin, which are thought to microphase separate *in vivo*, govern nuclear morphology, we implemented a composite coarse-grained polymer and elastic shell model. By varying chromatin volume fraction (density), heterochromatin levels and structure, and heterochromatin-lamina interactions, we show how the spatial organization of chromatin polymer phases within the nucleus could perturb nuclear shape in some scenarios. Increasing the volume fraction of chromatin in the cell nucleus stabilizes the nuclear lamina against large fluctuations. However, surprisingly, we find that increasing heterochromatin levels or heterochromatin-lamina interactions enhances nuclear shape fluctuations in our simulations by a “wetting”-like interaction. In contrast, shape fluctuations are largely insensitive to the internal structure of the heterochromatin, such as the presence or absence of chromatin-chromatin crosslinks. Therefore, our simulations suggest that heterochromatin accumulation at the nuclear periphery could perturb nuclear morphology in a nucleus or nuclear region that is sufficiently soft, while stabilization of the nucleus via heterochromatin likely occurs through mechanisms other than chromatin microphase organization.

## Introduction

The mammalian genome confined inside the ∼10 μm nucleus is composed of multiple extremely long DNA polymer chains, referred to as chromosomes. Chromosomes are packaged heterogeneously into chromatin fibers through structural and functional DNA-associating proteins. These chromatin fibers are dynamically organized into polymer phases^1–10^, generally known as euchromatin and heterochromatin. The spatial organization of chromatin is essential for regulating vital biological processes^10–13^, such as transcription, replication, and differentiation. Perturbations and alterations to chromatin organization are associated with a variety of human diseases and conditions, including aging, progeria, muscular dystrophy, and various cancers^14–19^. Recent experiments have also established that perturbed chromatin organization is causally connected to disruptions in nuclear shape^20–25^, which has long been used as a hallmark of these conditions. Moreover, recent research has shown that there are mechanistic links between alterations to nuclear architecture and perturbations to nuclear function, suggesting another intriguing link between chromatin spatial organization and nuclear function. Nonetheless, the physical mechanisms through which chromatin governs nuclear shape are not fully explained, and it is unclear whether and how the overall (micro)phase organization of chromatin contributes to nuclear morphology.

Chromatin has two main types, euchromatin and heterochromatin, which are distinguished by their transcriptional activities, compaction, and organization^8,26,27^. Euchromatin is transcriptionally active and maintains an open, accessible structure. Heterochromatin is composed of compacted chromatin domains in which transcriptional activity is low and chromatin is generally inaccessible. Regions of heterochromatin are further classified as facultative and constitutive, depending on their histone modifications^8,26^. In conventional nuclear chromatin organization, heterochromatin-rich regions of chromatin are predominantly located at the nuclear periphery, beneath the nuclear lamina, whereas the euchromatin is generally located in the nuclear interior^28^. Various measurement techniques, including whole-nucleus micromanipulation^21,29,30^ to nuclear elastography^31^ indicate that nuclear heterochromatin has a larger elastic modulus than euchromatin. In addition, the heterochromatin fiber seems to have a longer persistence length (i.e., distance below which a polymer behaves like a rod) than euchromatin^32^, and this difference may influence chromatin organization via entropic effects^33,34^. Additionally, heterochromatin is thought to exhibit stronger self-attraction than euchromatin, which may result in a mesoscale phase separation of chromatin within the nucleus^5,35,36^.

Chromatin phase separation, in turn, is thought to spatially organize chromatin within the confines of the nucleus, which may deform in response to this internal organization. Treating cells with drugs that inhibit enzymes that alter histone modifications (e.g., deacetylases, demethylases, and methyltransferases), can either induce or inhibit the formation of shape abnormalities known as nuclear “blebs”^21,37^. These deformations are chromatin-filled protrusions that frequently rupture and correlate with increased DNA damage^37,38^, as well as various diseases^18,25^. Inhibition of histone deacetylases or methyltransferases increases euchromatic histone marks and increases nuclear blebbing, while inhibition of demethylases increases heterochromatin content and rescues nuclear shape^21,37^. Similarly, the loss of heterochromatin protein 1α (HP1α), a chromatin-crosslinking protein that can induce heterochromatin phase separation^39,40^, induces nuclear bleb formation^41^. Despite these observations, it is unclear whether such alterations to chromatin phase organization are directly responsible for shape anomalies..

Beyond localized shape disruptions, the spatial distribution of heterochromatin and euchromatin may influence the shape of the entire nucleus. Immunostaining experiments in mouse rod cells have shown that reorganization and repositioning of heterochromatin to the nuclear interior coincide with a change from a prolate ellipsoidal shape to circular disc-like nuclear morphology^42^. Similarly, loss of peripheral heterochromatin in Hutchinson Gilford Progeria Syndrome (HGPS) patient cells^43,44^ and aging *C. elegans* cells^45^ leads to widespread wrinkling and folding of the nuclear envelope. Furthermore, in HGPS cells, these shape changes can be reversed by heterochromatin compaction^21,37^. Hence, experimental evidence suggests that the 3D chromatin organization within the nucleus can affect global nuclear morphology.

Chromatin is further organized by interactions with the nuclear envelope. Chromatin may be localized to the envelope (e.g., to the lamina and embedded proteins) via lamins and other chromatin-tethering proteins or histone modifications and histone-modifying enzymes^42,46–49^. Any breakdown in these interaction mechanisms can also affect the 3D chromatin organization and, in turn, nuclear morphology^20,46^. For example, the knockdown of a chromatin-tethering protein, PRR14, led to a wrinkled nuclear morphology with large undulations on the nuclear surface^50^. Furthermore, in *S. pombe*, knockouts of chromatin-tethering LEM domain proteins exhibited reduced increased nuclear deformations, apparently due to loss of chromatin interactions with the nuclear envelope^51^. Together, these experiments suggest that robust chromatin interactions with the nuclear periphery are important for the maintenance of regular nuclear morphology.

Previous polymer modeling studies suggested that chromatin phase separation, chromatin-lamina interactions, and chromatin volume fraction all can contribute to 3D genome organization^2–5,35,36,52–55^ . Within a rigid boundary, competition between heterochromatin-driven phase separation and adsorption of heterochromatin to the lamina can control the spatial distributions of chromatin, and its subtypes, heterochromatin and euchromatin^3–5^. However, it is unclear whether these effects would be coupled to the morphology of a deformable polymeric shell, such as the lamina. Elastic shell models confining simple homopolymers have shown that chromatin compaction/stiffness can aid in resisting nuclear deformations^29,41,56^. However, simulations of a similar model have suggested that chromatin-lamina interactions can destabilize nuclear shape in some scenarios^57^. Nevertheless, whether chromosomes, considered as multiblock copolymers that can undergo phase separation, may govern nuclear morphology have not been explored systematically.

To investigate the connection between chromatin organization and nuclear morphology, we use molecular dynamics (MD) to simulate heterochromatin and euchromatin phase separation within a polymeric shell representing the nuclear lamina. Focusing on a model in which the nuclear lamina is relatively soft, we find that heterochromatin’s self-interactions and interactions with the nuclear lamina can cooperatively contribute to nuclear shape fluctuations. Our simulations show that the volume fraction of the chromosome polymers determines the strength of the effects of heterochromatin phase separation on nuclear morphological fluctuations. In addition, the interaction affinity of heterochromatin to the nuclear shell drastically influences nuclear morphology, resulting in distorted nuclear shapes when the affinity is strong. Strikingly, our results differ from the conventional picture of chromatin-driven nuclear morphology, in which collapsed heterochromatin, especially at the nuclear periphery, stabilizes nuclear shape. Nonetheless, our results are consistent with some experimental scenarios and suggest a new mechanism by which chromatin phase organization may govern nuclear morphology. Overall, our findings suggest that the physics of polymer mixtures and phase separation can explain aspects of chromatin-related nuclear morphology alterations.

## Methods

### Molecular model of mammalian cell nucleus

We model the genome by 8 bead-spring copolymer chains. These chains are self-avoiding coarse-grained Kremer-Grest^58^ triblock copolymers in implicit solvent (Figure 1A). These block copolymers model chromosomes as composed of self-interacting compartments, which were extracted from high-throughput chromosome conformation capture (Hi-C) data from mouse rod cells^5^. Four of the eight block copolymers in our model are derived from chromosome 1, while the other four are derived from chromosome 2, similar to previous modeling^5^. Each block copolymer is composed of 6002 beads. 1000 beads at one end of each block polymer are assigned as constitutive heterochromatin, and the remaining 5000 beads represent genomic loci that can form A-or B-type compartments^5^, generally corresponding to euchromatin and heterochromatin. Two additional beads are added to each end of the polymer chains to represent the telomeres. Each bead in a polymer chain represents 40 kilobase pairs (kb) of chromatin, and each bead has size *b* = 1*σ*, where *σ* is the simulation unit length scale. This resolution allows the construction of genomic contact maps comparable to Hi-C experiments studying chromatin compartmentalization^5,59–61^. In the simulations, the polymers are initially placed at random positions in a compact form^62^ near the center of the spherical volume within the shell (Figure 1A). Adjacent polymer beads are connected by non-extensible bond potentials, and bond crossings are disallowed (See Supplementary Information for further details of the polymer and shell model).

**Figure 1:**
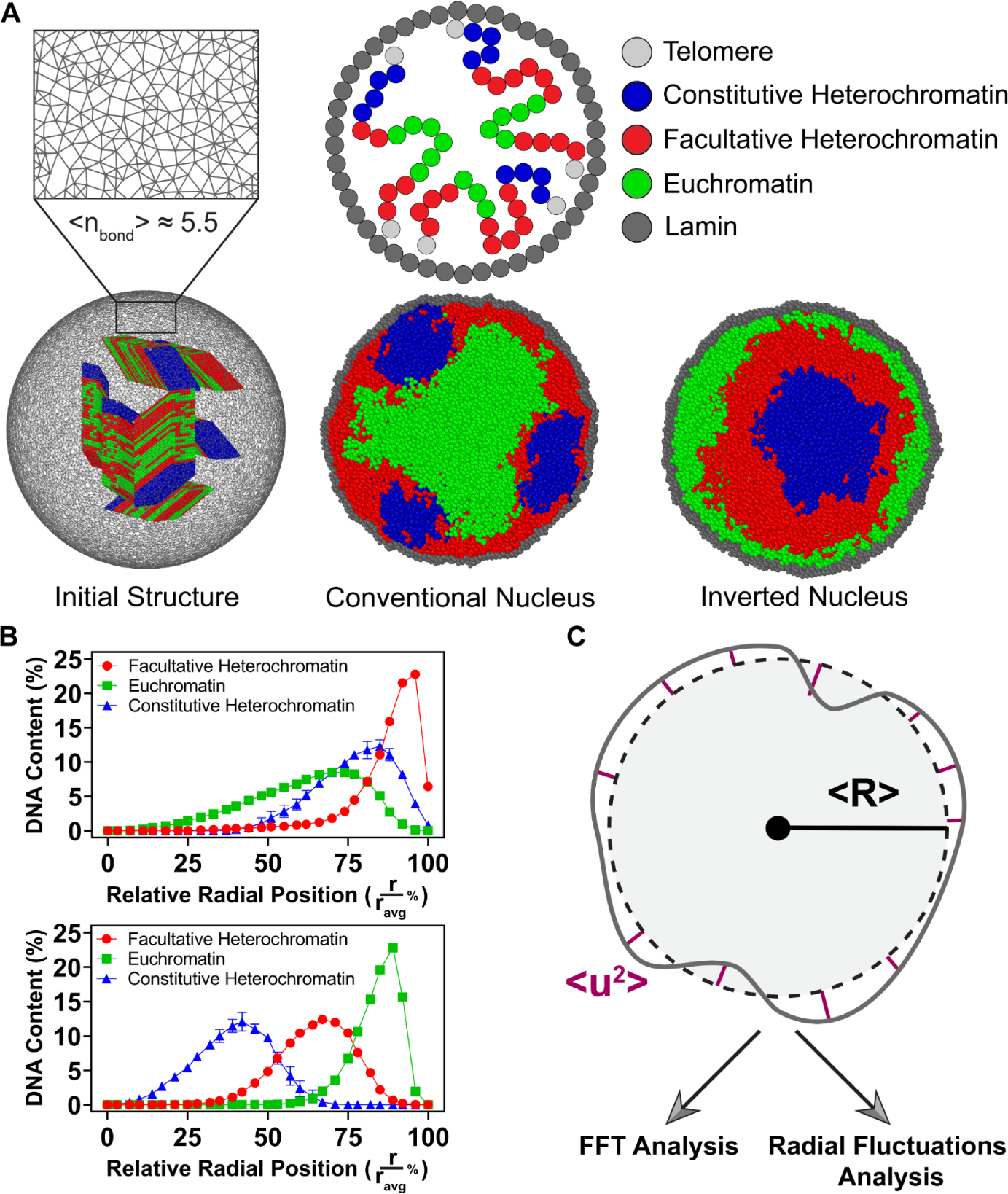
Initialization, equilibration, and analysis of the polymer and shell model for the cell nucleus. A) Schematic illustration of the MD simulation model, with block copolymer and deformable shell structures. Chromosome-like structures composed of 8 tri-block polymers are placed randomly and confined inside the shell. Initial structures were run sequentially to obtain conventional and inverted nuclei. B) The density profiles of the different polymeric blocks in our simulations in conventional and inverted nuclei after relaxation. C) A schematic illustration of the analysis and metrics used to quantify nuclear shape fluctuations.

Following previous modeling^29,56^, the beads composing the confining shell model the elasticity of the nucleus arising due to the nuclear envelope, including the lamina, a meshwork of lamin protein^63^. In the initial preparation of the shell, each shell bead is randomly placed on the surface of a sphere of radius *R*_0_=42*σ*. The beads are connected by non-extensible bond potentials. On average, each shell bead forms *n*_bond_ bonds to neighboring shell beads, with 5 ≤ *n*_bond_ ≤ 8. These bonds are initially assigned by a predefined distance parameter such that they bond with nearest neighbors and prevent void spaces on the shell surface (Figure 1A). Throughout the simulation, the total numbers of bonds and beads are constant.

### MD simulation parameters

Steric interactions between beads, whether they are bonded to each other or not, are modeled by a eLennard-Jones (LJ) potential with various cut-off distances (see the Supplementary text for simulation parameters). All simulations are run using the LAMMPS MD package^64^. The simulation timestep is Δt = 0.005*τ*, where *τ* is the unit time scale in the simulations. Initially, the simulations are run for 3×10^4^ MD steps to allow for the initial relaxation of the shell. During this phase of the simulations, the shell radius is shrunk by ∼30% as the assigned bonds reach their equilibrium length. The purpose of these simulations is to locally equilibrate the shell structure and reduce holes on the shell surface that could lead to chromatin escaping from the nucleus. Subsequently, an additional 10^5^ MD steps are run to relax the polymer chains and corresponding multiblock structures (Figure 1A). During the relaxation simulations, the attractive interactions between heterochromatin beads (i.e., constitutive and facultative heterochromatin and telomeres) and shell beads are switched on. We then perform data production simulations to obtain conventional nuclear organization with the interaction parameters listed in the Supplementary Information. The conventional nucleus simulations are run for 5×10^6^ MD timesteps to obtain a quasi-equilibrium chromatin distribution (Figure 1B). After recapitulating conventional nuclei, we turned off the interactions between heterochromatic polymer subunits and the shell to recapitulate inverted nuclei^5^. The inverted nucleus simulations are run for 10^7^ MD timesteps. The number of simulation steps for equilibration in conventional and inverted nucleus simulations was determined from chromatin radial distribution and density plots (Figure 1B & Supplementary Figure S1). The polymer volume fraction *φ* is varied by decreasing the initial number of polymer blocks from *n_block_*=8 to *n_block_*=6 and *n_block_*=4 to obtain *φ*=19% and *φ*=14%, respectively, in addition to the default value of *φ*=23%.

### Calculation of nuclear shape fluctuations

To calculate nuclear shape fluctuations, we computed the Fourier modes^56,65,66^ by the Fast Fourier Transform algorithm (FFT)^67^. At each timestep, we considered a thin slice of the shell with a width of *w=*1σ. For each bead in the slice, we calculated its displacement from the average radius of the slice (Figure 1C). We run the FFT algorithm on the obtained radial distances for the lowest 230 modes, *q*, and take the time average of the fluctuation amplitudes over the last half of the simulation. We also measured the time-averaged standard deviation of the real-space displacement from the mean radial position to further quantify shape fluctuations.

## Results

### High Polymer Volume Fraction Suppresses Nuclear Shape Fluctuations

Recent studies suggest that chromatin density might influence the size and shape of the nucleus via polymer entropic pressure from chromatin^68–70^. In parallel, chromatin decompaction can increase nuclear size^71^ and induce chromatin-filled protrusions^23^. Conversely, hyperosmotic shock alters nuclear architecture by collapsing chromatin^72–75^.

Therefore, to understand the interplay between chromatin organization and nuclear shape, we performed simulations in which we varied the volume fraction of chromatin polymers confined inside the elastic, deformable shell representing the lamina (see Methods). In our simulations, we considered two scenarios for nuclear organization: 1) conventional organization with heterochromatin at the periphery and euchromatin in the interior and 2) “inverted” organization with heterochromatin in the interior and euchromatin at the periphery (Figure 2)^5,42^. The conventional chromatin distribution models many cells in which heterochromatin is preferentially located near the nuclear periphery, while the inverted organization models mouse rod cell^5,42^ and serves as the limiting case in which peripheral heterochromatin levels vanish (Figure 1A).

**Figure 2:**
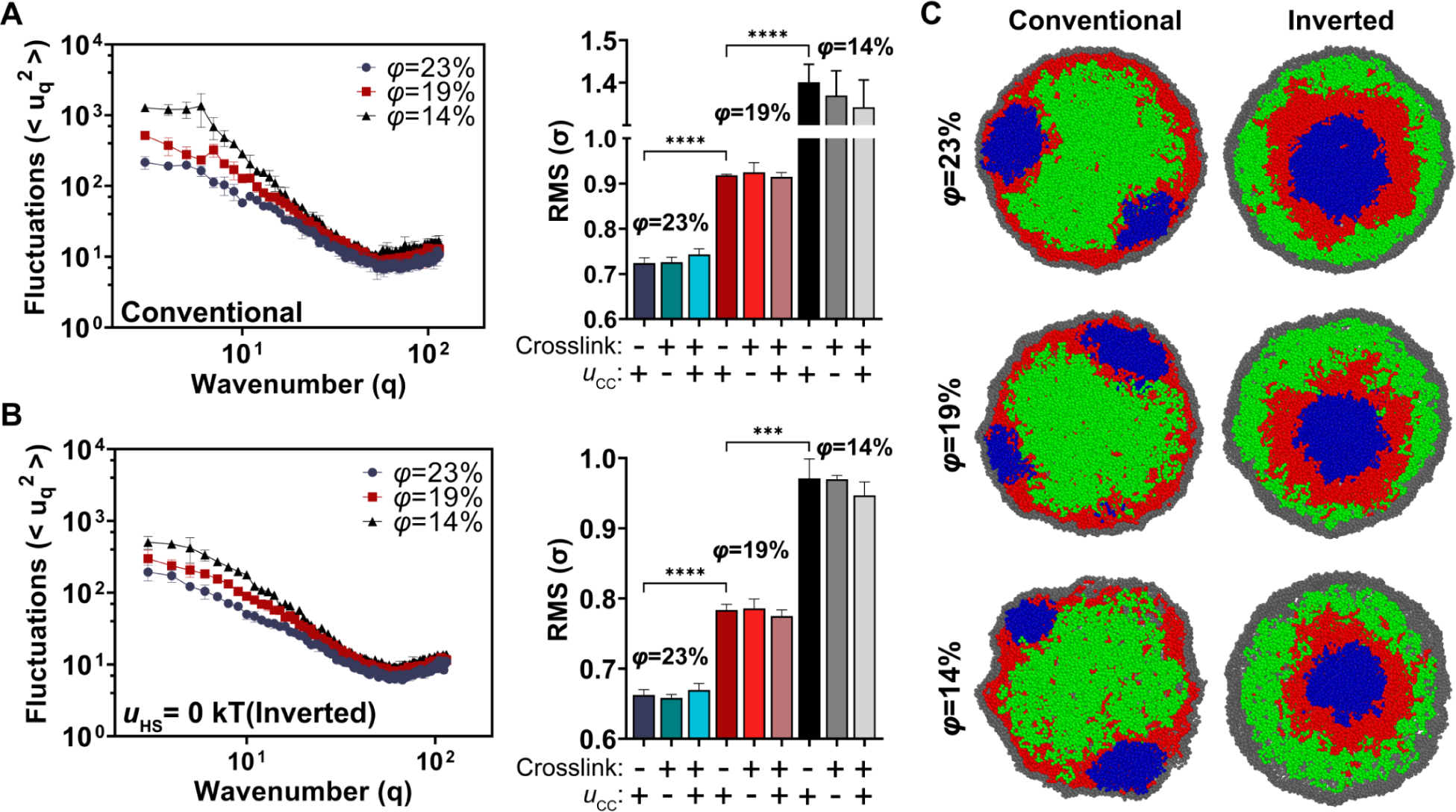
The effect of polymer volume fraction on nuclear shape fluctuations in conventional and inverted nuclei. A) Shape fluctuation analysis was conducted on simulations at various volume fractions for conventional nuclei at three different volume fractions (φ=14%, φ=19%, and φ=23%). The constitutive heterochromatin domains (blue) are crosslinked or not and the LJ potential, u_CC_, between constitutive heterochromatin beads, is switched on and off, as indicated by +/-beneath each bar in the bar plot at right. B) The Fourier spectrum of fluctuations and the RMS amplitudes plotted for inverted simulations. C) Representative snapshots of chromatin spatial organization and nuclear morphology in conventional and inverted scenarios for different chromatin volume fractions

In simulations, we characterized nuclear shape fluctuations by calculating the deviations of the simulated structure from a sphere. We computed the Fourier spectrum of fluctuations as well as the root-mean-square (RMS) deviation of the elastic shell radius from a sphere with a constant mean radius^45,51,76^ (see Methods). Our analyses show that decreasing the chromatin volume fraction increases the amplitudes of nuclear shape fluctuations, particularly for modes with small wavenumbers, *q*, corresponding to length scales on the order of nuclear radius (Figure 2A). Thus, higher chromatin volume fraction globally stabilizes nuclear morphology, likely due to increased polymer osmotic pressure. In inverted nuclei, we observe a similar effect of chromatin density (Figure 2B,C), albeit with weaker fluctuation amplitudes as compared to the conventional organization in the low *q* limit (i.e., nuclear-scale fluctuations).

### Nuclear Shape Fluctuations are Insensitive to the Internal Structure of Heterochromatin

Across all volume fractions, we observed that the nucleus with conventional chromatin organization exhibits systematically higher fluctuation amplitudes and RMS shape fluctuations as compared to inverted nuclei (i.e., 40% higher fluctuations at the lowest volume fraction, *φ*=14%,) (Figure 2A,C). This contrasts with the idea that peripheral heterochromatin stabilizes nuclear morphology, so we next investigated whether these effects could be modulated by altering the internal structure of heterochromatin, which may be facilitated *in vivo* by alterations to histone modifications^7,75,77–79^ or chromatin-binding proteins, such as HP1^9,39,40^.

Our standard chromatin model consisted of constitutive heterochromatin beads that had nonspecific attractive interactions with each other. In addition to this model, we tested whether intra-domain linkages between constitutive heterochromatin beads (i.e., chromatin-chromatin crosslinks) could alter nuclear shape fluctuations. For the crosslink model, constitutive heterochromatin beads were permanently crosslinked to each other by bonds similar to those connecting polymer chains.

We performed simulations at each of the three different volume fractions with and without constitutive heterochromatin crosslinking and with or without nonspecific attractive constitutive heterochromatin to constitutive heterochromatin attractions (Figure 2A,B). Simulations showed that the absence of crosslinking influences heterochromatin spatial organization in conventional nuclei; constitutive heterochromatin domains tended to merge (Supplementary Figure S3). However, the distribution of constitutive heterochromatin domains did not alter the shape fluctuations, as measured by either the Fourier spectrum or the RMS shape fluctuations of the shell (Figure 2). The presence or absence of crosslinking also did not change shape fluctuations in inverted nuclei. Similarly, we observed that shape fluctuations were insensitive to the presence or absence of nonspecific heterochromatin self-attractions, provided that intra-domain crosslinks were present. Altogether, these results indicate that heterochromatin’s spatial positioning, but not necessarily its internal structure, is critical in determining the amplitudes of shape fluctuations of the confining elastic shell.

### Interactions between Heterochromatin and Lamina Alter Nuclear Shape Fluctuations

To further understand the mechanisms by which peripheral heterochromatin may govern nuclear shape, we altered the interactions between heterochromatin and the lamina in our simulations. Experiments have shown that heterochromatin-lamina interactions are critical in regulating nuclear morphology^50,80–82^. We modeled these interactions through a nonspecific (nonbonded) attractive interaction with strength *u*_HS_ between heterochromatin and lamina (shell) beads. By varying *u*_HS_, we investigated how heterochromatin-lamina interactions may regulate nuclear shape fluctuations, emulating variations in levels or interactions with chromatin-lamina tethering proteins, such as LBR^42,83,84^.

Our analysis reveals that increasing the strength of the heterochromatin-shell attraction significantly increases nuclear shape fluctuations regardless of the chromatin volume fraction (Figure 3A-C and Supplementary Figure S4). For nuclei with strong heterochromatin-lamina attractions, we observe large bulges and localized wrinkles on the shell (right column of Figure 3C). With strong interaction energies, although the radial pattern of chromatin organization is preserved, each heterochromatin domain distort the shell by wrapping it around the surface of the domain (Figure 3C). At the strongest attraction energy studied (*u*_HS_=3 kT, where 1 kT ≈ 0.6 kcal/mol), shape fluctuations increase drastically across all length scales (Figure 3A). Decreasing the attraction affinity from our standard value, *u*_HS_=0.75 kT, to a lower value, *u*_HS_=0.375 kT, reduces nuclear shape fluctuations, particularly in the small-wavenumber (large lengthscale) regime (Figure 3A).

**Figure 3:**
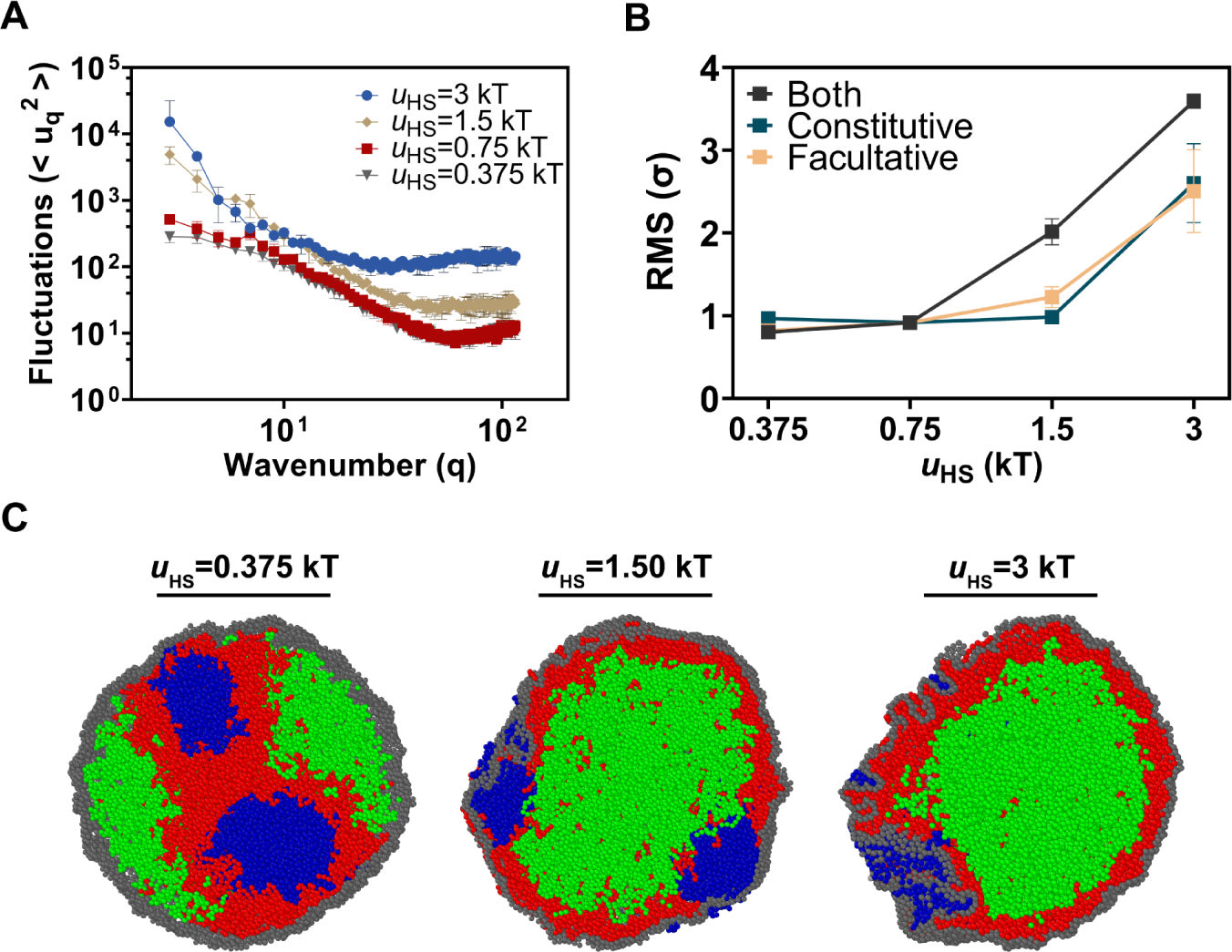
Heterochromatin-shell interactions influence nuclear shape fluctuations and morphology. A) Nuclear shape fluctuations were calculated in conventional nucleus simulations in which the interaction between heterochromatin (facultative and constitutive) and the shell was altered. The volume fraction was set to φ=19%. B) RMS nuclear shape fluctuations in three different scenarios of heterochromatin-lamina interactions: one in which only constitutive heterochromatin is attracted to the lamina, one in which only facultative heterochromatin is attracted to the lamina, and one in which both types of heterochromatin are attracted to the lamina. C) Snapshots of three different simulations with different strengths of heterochromatin-lamina interactions., where both heterochromatin types are attracted to the lamina.

However, in this scenario, the spatial organization of chromatin is altered. Facultative heterochromatin near the nuclear periphery is reduced, forming mesoscale, phase-separating heterochromatin domains (Figure 3C). When heterochromatin-shell interactions are set to *u*_HS_= 0, which is the case of inverted nuclei, morphological fluctuations on even the smallest length scales (largest *q*) become much weaker (Supplementary Figure S4). The finding that chromatin-shell interactions can enhance nuclear shape fluctuations ran counter to our expectation that such interactions would stabilize nuclear shape, as they frequently do *in vivo*^20,50,51^. We therefore tested whether this phenomenon is sensitive to the stiffness of the shell. In simulations with stiffer bonds between shell beads (effectively, a higher bending rigidity^56^), we found that the destabilizing effect of heterochromatin-shell interactions is somewhat weakened for the volume fractions we considered here. Altogether, our results suggest that attractive heterochromatin-shell interactions can enhance nuclear shape fluctuations under conditions with a sufficiently soft confining shell.

Since the chromatin polymer in our simulations is composed of two types of heterochromatin, constitutive and facultative, we also separately vary their interactions with the confining polymeric shell (Figure 3B). We find that increasing the interaction strength of either type of chromatin with the lamina is sufficient to influence the nuclear shape fluctuations (Figure 3B). Nevertheless, increasing only the facultative heterochromatin interactions with the shell causes the facultative heterochromatin to coat the inner surface of the shell completely, replacing the more weakly interacting constitutive heterochromatin. The same mechanism does not appear with the constitutive heterochromatin due to strong heterochromatin-heterochromatin interactions collapsing these chromatin domains (Supplementary Figure S8).

On top of these changes, nuclear morphology for each perturbation is governed by the chromatin volume fraction. Strong shape fluctuations are further strengthened by lower chromatin volume fractions (Supplementary Figure S4-S9). Together, these findings indicate that the nuclear shape fluctuations can be affected by both heterochromatin types but via different mechanisms in the model.

### Heterochromatin Self-affinity Alters its Spatial Distribution together with Nuclear Shape Fluctuations

Since changes to heterochromatin-lamina interactions alter the spatial distribution of heterochromatin, we sought a complementary method to probe whether and how the spatial distribution of heterochromatin can perturb nuclear shape fluctuations. Therefore, we studied the effects of changing the self-affinity of heterochromatic regions, as might occur when alterations are made to histone modifications^75,77^.

Our simulations show that the strength of heterochromatin self-affinity can alter both the spatial distribution of chromatin and nuclear shape fluctuations. This is exemplified in Figure 4, where from left to right, heterochromatin-heterochromatin attractions increase, and from top to bottom heterochromatin-shell interactions increase. Abolishing the self-affinity of the facultative heterochromatin monomers (i.e., *u*_HH_=0, left column of snapshots in Figure 4) decreases peripheral heterochromatin levels. Facultative heterochromatin beads coat the interior of the lamina, but additional heterochromatin has no preference for the nuclear periphery, and thus, disperses throughout the nuclear interior.

**Figure 4.**
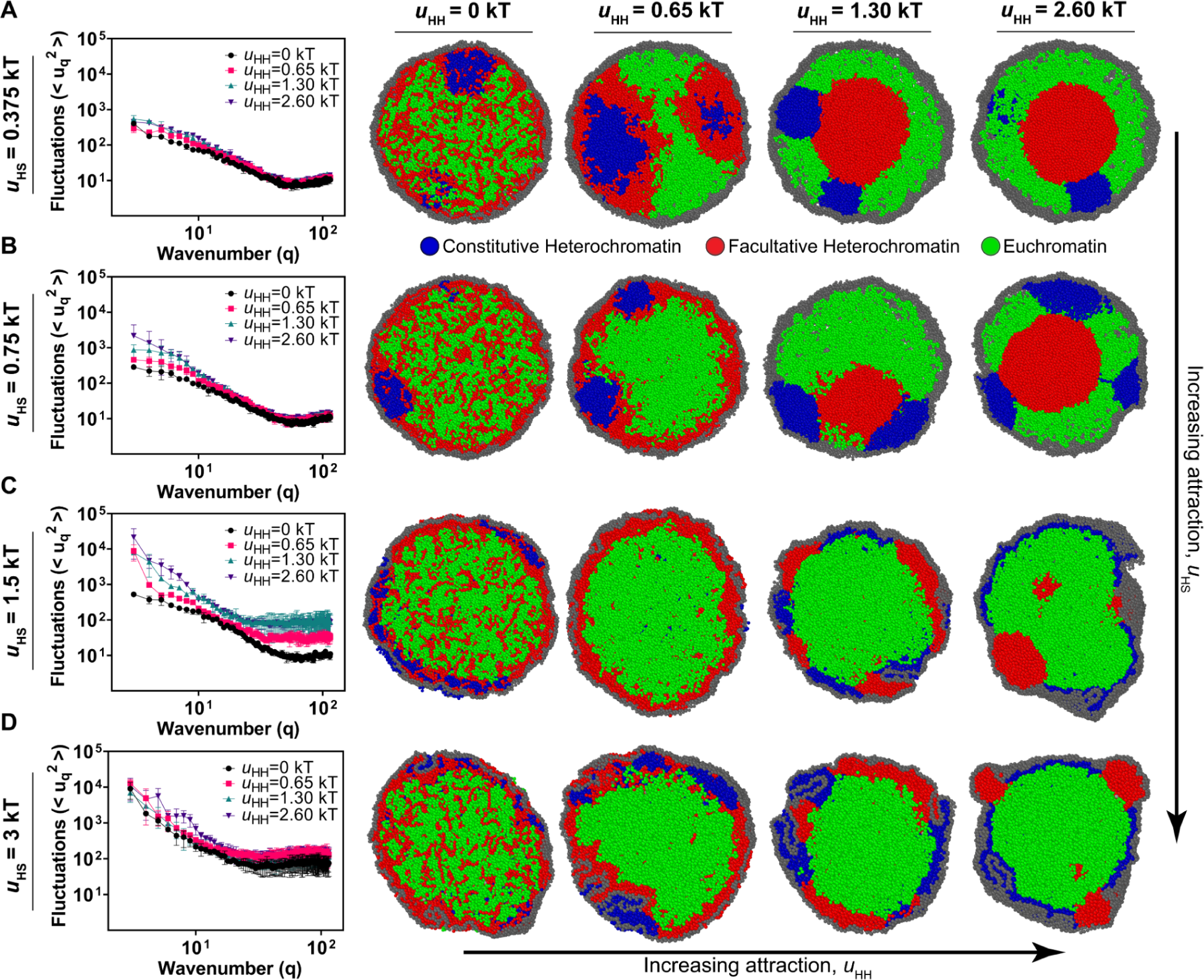
The interplay between heterochromatin-heterochromatin and heterochromatin-shell interactions. A-D) Fourier spectra of nuclear shape fluctuations and corresponding snapshots for different self-affinities of heterochromatin (different for different colors in each plot and increasing from left to right across each row) and different heterochromatin-shell interactions strengths (increasing from top row to bottom row). A) The attraction affinities between facultative heterochromatin domains are varied, and the shell is set to u_HS_=0.375 kT. The polymer volume fraction is set to φ=19%. B) The attraction affinity between shell and heterochromatin domains is set to u_HS_=0.75 kT. C) The attraction affinity between shell and heterochromatin domains is set to u_HS_=1.5 kT. D) The attraction affinity between shell and heterochromatin domains is set to u_HS_=3 kT.

Peripheral organization of heterochromatin can also be disrupted by increasing the self-affinity of facultative heterochromatin to values larger than the chromatin-lamina attraction strength, i.e., *u*_HH_ > *u*_HS_ > 0 (Figure 4A-B, right). In this case, facultative heterochromatin forms a phase-separated region at the center of the nucleus, much like an inverted nucleus, as observed in a previous study^5^. Constitutive heterochromatin, in contrast, remains at the periphery since its self-affinity is unchanged in these simulations. Interestingly, this large-scale reorganization of heterochromatin positioning does not dramatically alter the shape fluctuations when chromatin-lamina interactions are weak (Figure 4, top two rows, e.g., compare *u*_HH_=0.65 *kT* vs. *u*_HH_=1.30 *kT*). Thus, there is only a modest difference between the shape fluctuations of conventional and inverted nuclei in this scenario.

Since we observe stronger shape fluctuations in simulations with strong chromatin-lamina interactions, we hypothesized that interactions with the densely packed peripheral heterochromatin layer beneath the shell surface could drive shape fluctuations (Figure 4A). Increasing the chromatin-lamina attraction strength leads to strong adsorption of heterochromatin to the interior of the lamina shell (Figure 4B-D). With strong chromatin-lamina interactions, even when the heterochromatin self-affinity is nonexistent (*u*_HH_=0), a peripheral heterochromatin layer forms near the shell’s interior surface and distorts the shell, increasing nuclear shape fluctuations (Figure 4D, leftmost image). Interestingly, if the heterochromatin-shell attraction is nonzero, heterochromatin further accumulates near the periphery. In turn, this heterochromatin accumulation can further increase nuclear shape fluctuations (Figure 4B-D). These results indicate that heterochromatin-lamina interactions can facilitate chromatin polymer phase separation and that heterochromatin self-affinity and lamina interactions can cooperatively increase nuclear fluctuations.

Increasing the heterochromatin-shell attraction affinity to large values (e.g., *u*_HS_ > *u*_HH_) leads to the localization of all heterochromatin at the periphery, depleting heterochromatin from the nuclear interior (Figure 4C, D). Nevertheless, the uniformity of the peripheral facultative and constitutive heterochromatin phases is highly disrupted. In this case, constitutive heterochromatin is mixed with facultative heterochromatin due to the strong attraction of all heterochromatin with the shell overwhelming the self-affinity of different heterochromatin phases (snapshots on the right side of Figure 4C, D). Notably, the shape deviations from a sphere are elevated as the heterochromatin-shell interactions are further increased (Figure 4C,D).

In the case where heterochromatin-shell and heterochromatin-heterochromatin interactions are both strong, a new nuclear morphology emerges. Spherical domains of facultative heterochromatin at the periphery create chromatin-filled protrusions that bulge outward from the shell surface (Figure 4D, right column). While these structures are visually reminiscent of experimentally observed nuclear bleb^21,23,37,43,85^, the simulated “blebs” are filled with heterochromatin, unlike the blebs observed in a variety of previous experiments, which generally contain less compact euchromatin^21,63,86,87^. Therefore, heterochromatin self-affinity and spatial distribution can alter nuclear morphology in our simulations. Furthermore, our model predicts a new polymer-physical mechanism through which bleb-like protrusions can be formed in cell nuclei with soft shells.

### Nuclear Heterochromatin Content Affects Shell Morphology and Fluctuations

Studies show that decondensing chromatin, decreasing the overall level of heterochromatin or increasing the total amount of euchromatin is associated with abnormal nuclear morphology^21,23,37^. We therefore investigated whether the fraction of facultative heterochromatin per chromosome alters nuclear shape fluctuations in our model.

To model these changes, we randomly converted euchromatin beads into facultative heterochromatin beads or vice versa. We varied the percentage of the chromatin fiber composed of facultative heterochromatin fraction from *f=*10% to *f=*75% (compared to our standard value of *f* ≈ 50%).

We find that increasing the facultative heterochromatin content elevates nuclear shape fluctuations and introduces morphological abnormalities (Figure 5A). Up to *f=*55%, primarily lower mode (larger lengthscale) fluctuations are elevated (Figure 5B). However, as the heterochromatin fraction increased toward *f=*75%, fluctuations increased across all wavenumbers. This manifested as a distorted nuclear morphology with wrinkles, bulges, and other features at various length scales (Figure 5C). The effect of higher facultative heterochromatin is enhanced at lower volume fractions and weakens as the volume fraction is increased (Figure 5B). Turning off the heterochromatin-shell attractive interactions (i.e., inverted nucleus) reduces the shape fluctuations in the large *f* limit, suggesting that the observed effects are once again due to peripheral heterochromatin and its interactions with the lamina (Supplementary Figure S16-19).

**Figure 5).**
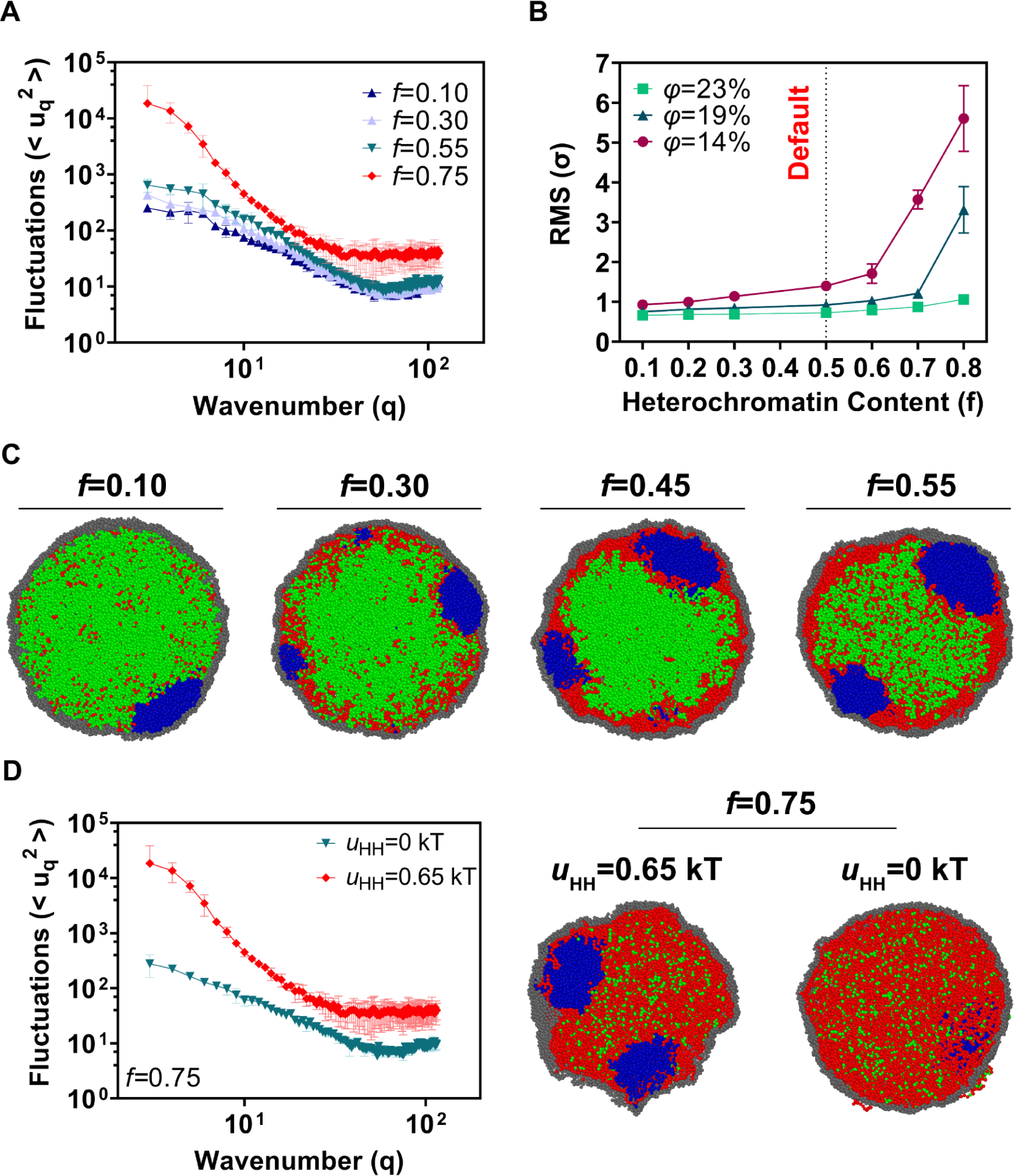
The facultative heterochromatin content at the nuclear periphery impacts the nuclear morphology. A) Fourier spectra of shape fluctuations for conventional and inverted nuclei simulations, where the facultative heterochromatin content is set to f=10%, f=30%, f=45%, f=55%, and f=75%. The polymer volume fraction is set to φ=19%. B) RMS fluctuation amplitudes for varying levels of facultative heterochromatin at different volume fractions, with the value for standard simulations labeled as “default.” C) Representative simulation snapshots for simulations with different levels of facultative heterochromatin. D) Shape fluctuations with and without heterochromatin self-affinity for f=75% case, along with simulation snapshots. Reducing self-affinity inhibits shape fluctuations in this scenario.

Nevertheless, increasing heterochromatin levels in experiments generally leads to a more rounded nuclear shapes without nuclear blebs and shape disruptions^21,37,88^, which contradicts our simulation results (Figure 5C). We speculated that the combined effects of facultative heterochromatin self-attraction and attraction to the shell could be destabilizing nuclear shape in our simulations. We therefore repeated the high-heterochromatin simulations (i.e., *f=*75%) without heterochromatin self-affinity at the lowest volume fraction, for which we had observed the most drastic shape anomalies (i.e., *φ*=14%). We then observed decreased shape fluctuations and less deformed morphologies (Figure 5D). However, as before (Figure 4), while weaker heterochromatin self-attractions decreased shape fluctuations, they also inhibited facultative heterochromatin phase separation in the nuclear interior and, albeit to a lesser extent, at the periphery (Supplementary Figure S11-12). Overall, our simulations showed that increasing the heterochromatin levels elevated nuclear shape fluctuations due to widespread distortions of the lamina via heterochromatin-lamina attractions, but heterochromatin decondensation was able to suppress this effect.

## Discussion

Using coarse-grained MD simulations, we modeled isolated cell nuclei encapsulating chromosomes to study the effects of heterochromatin organization on nuclear morphology. Chromosomes, which are composed of euchromatin and facultative and constitutive heterochromatin, were modeled as block copolymers containing three types of monomeric subunits^5^. We used an elastic polymeric shell model of the nuclear lamina to sterically confine the polymers within the nuclear volume^29,56,57^. Our model produces a broad spectrum of chromatin spatial distributions and nuclear shapes, depending on chromatin volume fraction, heterochromatin self-affinity and internal structure, heterochromatin-lamina interactions, and total heterochromatin content. We discuss these findings in detail below.

### Chromatin volume fraction can repress nuclear shape fluctuations

Our simulations suggest that the volume fraction of the chromatin polymers can regulate the nuclear shape fluctuations and morphology. In particular, a higher volume fraction of chromatin supports a more regular nuclear shape (Figure 2), consistent with previous experiments and simulations in which chromatin was degraded (experiments) or not present (simulations) in the nucleus^56^. In our simulations, this is largely due to increased polymer osmotic pressure exerted outward on the nuclear lamina, which swells the nucleus and inhibits fluctuations. Consistently, in simulations higher in collapsed heterochromatin, polymer-based osmotic pressure (i.e., translational entropy) is limited, and shape fluctuations increase (Figure 4A-C, Figure 5). In simulations, pressure due to the internal polymer is insufficient to generate localized nuclear deformations (i.e., blebs), but our observations are consistent with nuclear blebs *in vivo* generally containing transcriptionally active euchromatin^21,63,86,87^.

Experimentally, hyperosmotic osmotic pressures can lead to rapid chromatin collapse^72–74,89,90^. Although this condition corresponds to a shrinking nucleus and a correspondingly higher chromatin mass density, chromatin itself collapses due to the change in salt conditions. This, in turn, reduces chromatin polymeric pressure. Consistently, hyperosmotic shock induces morphological distortions on the nuclear surface, similar to what we observe in our simulations with low chromatin volume fractions (Figure 2). Inversely, hypoosmotic shock leads to chromatin swelling, and consequently, more spherical nuclear shape^91^, much as we observe in simulations with larger chromatin volume fractions. Therefore, we conclude that the effective volume fraction of chromatin within the nucleus can directly impact nuclear morphology.

### Heterochromatin near the nuclear periphery can disrupt morphology of soft nuclei

Surprisingly, we found that compaction and overall levels of heterochromatin, particularly at the nuclear periphery, can drive large and widespread nuclear shape fluctuations across multiple lengthscales. This runs counter to experiments showing that high levels of heterochromatin^21,37^, linkages within heterochromatin^41^, and interactions between chromatin and the nuclear envelope all generally^20,50,51^ stabilize cell nuclear shape. Nonetheless, our simulations mirror findings in several specific scenarios with abnormal heterochromatin or lamina-associated proteins.

In particular, in progeria patient cells bearing the lamin A/C mutation E145K, heterochromatin is enriched at regions of the nuclear envelope that exhibit large bleb-like protrusion^44^. These distorted regions are further notable for being depleted of lamin A/C, meaning that they are likely less stiff than the rest of the nuclear lamina^29,92–94^. Thus, in this scenario, the dense heterochromatin phase localized to the nuclear periphery may drive large nuclear shape aberrations by overcoming the elastic bending energy of the lamina through the combination of phase and adsorption interactions.

Our simulations predict that elevated binding of heterochromatin to the nuclear periphery can also promote nuclear morphological distortions. This contrasts with several experiments that depleted chromatin-binding proteins mediating interactions with the lamina^50,51,95^. Nonetheless, our simulations are consistent with other studies, such as a report in which cells inducibly expressing lamin A mutant progerin formed nuclear blebs and other abnormal nuclear morphologies^57^. Progerin-expressing cells also exhibited more frequent interactions between the lamina and polycomb heterochromatin. Corresponding simulations suggested that these interactions could be responsible for disrupting nuclear morphology.

Beyond this report, our results also appear to be consistent with experiments in which the lamin B receptor (LBR) expression levels are altered^84^. In differentiating skin cells, nuclei appear to present abnormal morphology when LBR is overexpressed^83^. Similarly, for neutrophils, it was shown that LBR expression is essential for obtaining highly lobulated non-spherical nuclei that are typical of those cells^84,96^. Intriguingly, depletion of LBR in mouse neutrophils is associated with a redistribution of heterochromatin to the nuclear interior^97,98^, similar to more recent observations in thymocytes^5^. Our simulations indicate that these experimental observations collectively could be due to strong attractive interactions between heterochromatin and the nuclear lamina.

Such interactions can disrupt nuclear shape by bending the lamina to maximize interactions with the heterochromatin phase (Figure 4). Strong intra-heterochromatin interactions that compact heterochromatin could potentially amplify these effects (Figure 4). This would be especially notable in nuclei in which heterochromatin forms globular foci rather than a compact layer at the periphery. Further simulations showed that these effects are relatively weaker in simulations of stiffer nuclei (Supplementary Figure S21), suggesting that heterochromatin phase separation could be disruptive particularly in nuclei or regions of nuclei that are more liquid-like or depleted of lamins. Consistent with this hypothesis, lamin A/C levels are low in granulocytes such as neutrophils^99,100^ where LBR expression deforms nuclei, while they are higher in fibroblasts in which LBR *depletion* leads to nuclear shape perturbations^101^. Thus, we predict that in some scenarios, heterochromatin compaction into mesoscale phases and heterochromatin-shell interactions could generate aberrant nuclear shape deformations.

### Other possible mechanisms for regulation of nuclear shape by heterochromatin

To the extent that heterochromatin phase separation may enhance nuclear shape fluctuations and aberrations *in vivo*, our coarse-grained simulations suggest that the phase organization of heterochromatin does not generically stabilize the nuclear envelope against fluctuations. Therefore, the mechanistic question of how heterochromatin stabilizes cell nuclei against deformations, blebbing, and rupture remains open.

Our simulations modeled an isolated nucleus with a relatively soft lamina in a quasi-equilibrium environment. Consequently, the simulated nuclei were not subject to external stresses due to intracellular and extracellular forces. Various studies have indicated that interactions between the nucleus and the cytoskeleton can drive, and in some cases, suppress changes to nuclear morphology in a variety of scenarios^51,65,76,102–105^. In cases where heterochromatin or chromatin condensation inhibits alterations to nuclear shape^21,24,29,37,41,106,107^, it may act by increasing nuclear stiffness to resist mechanical forces applied to the nucleus^29,41,107^. Beyond external driving, it remains unclear to what extent heterochromatin might stabilize nuclear morphology against internal perturbations, such as nonequilibrium driving by transcription and other motor activities^66,87^. Therefore, while chromatin condensation can have a disruptive effect on isolated nuclei, there are likely other mechanisms underpinning heterochromatin’s effects on nuclear morphology *in vivo*.

### Physical mechanism of heterochromatin-induced morphological fluctuations

In our simulations, increased heterochromatin compaction and levels increased nuclear morphological fluctuations (Figures 3 and 5). This effect may arise by one of several physical mechanisms. Increasing the amount of heterochromatin or its self-affinity increases the amount of collapsed polymer in our simulations. In polymer physical terms, increasing heterochromatin increases the amount of polymer in poor solvent conditions. This generates a lower polymeric pressure inside the nucleus, which allows the lamina, a polymeric shell (or solid membrane) to fluctuate and distort. Consistently, simulations without heterochromatin self-affinity leads to smaller shape fluctuations due to increased polymeric pressure (which is due to increased translational entropy of heterochromatic regions), even at the highest levels of heterochromatin (Figure 5D). These observations parallel the changes in nuclear morphology that we observe when decreasing the chromatin polymer volume fraction (Figure 2).

Complementarily, polymer self-affinity combined with attractive interactions to a peripheral boundary can physically deform the boundary to lower the free energy of the system. Experiments with proteins condensing within lipid vesicles showed that strong phase separation can lead to the formation of long protein-coated lipid tubules and other structures, thus creating various vesicle morphologies^108–110^. Similarly, theoretical modeling has indicated that phase condensation of polymer solutions on a biopolymeric surface can generate morphological deformations on the surface^111,112^. More broadly, several experiments have recently demonstrated that biological phase separation can induce forces on substrates ranging from single-molecule DNA to chromatin^113–115^. Such scenarios arise due to wetting of membranes (a “solid membrane” or shell in our case) by biopolymer-rich phases, resulting in competition between substrate stiffness and surface tension of the phase-separated domains. In our simulations, we observed such wetting-induced-like morphological transitions (blebs and wrinkles) when heterochromatin-shell interactions are strong relative to heterochromatin self-affinity (Figure 4). That is, if heterochromatin’s tendency to form an interface with euchromatin dominates over the tendency to form an interface with the shell, inverted nuclear organization can emerge. However, the opposite case of stronger heterochromatin-shell interactions can result in highly distorted nuclear morphologies, provided that the shell is soft enough (e.g., comparable to the bending stiffness of lipid membranes).

Furthermore, while regular copolymers (i.e., with well-defined blocks) are known to undergo microscopic phase separation and exhibit highly regular phase^116^, random copolymers can form heterogeneous local domains^117^. However, the mechanisms explored by our simulations require nuclear-scale alterations to heterochromatin phase organization to generate nuclear-scale shape aberrations. We observed that reorganization may occur by small localized domains merging with larger mesoscopic domains near the periphery. This increases the nonuniformity of chromatin organization at the nuclear periphery, which facilitates larger effects on nuclear shape. Thus, epigenetic changes that promote large-scale phase formations at specific peripheral sites are most likely to disrupt nuclear shape.

In summary, our model suggests that chromatin organization, via chromatin volume density and heterochromatic phase separation, can alter nuclear morphology. Strikingly, heterochromatin phase separation, especially at the nuclear periphery, *promotes* disruption of nuclear shape. Since heterochromatin is generally considered to stabilize nuclear shape, our simulations suggest that the architectural effects of heterochromatin phase separation must be balanced by other aspects of nuclear mechanical response and organization. However, our model predicts that in nuclei with soft laminas or excessive heterochromatin recruitment to the nuclear lamina, this balance may be disrupted to form blebbed or lobulated nuclei, as it appears to be in granulocytes and some disease scenarios. Therefore, our simulations predict a new, largely unappreciated mechanism by which chromatin may regulate nuclear morphology.

## Acknowledgements

We thank Martin Falk for instructive discussions. EJB thanks Andrew Stephens and John F. Marko for helpful discussions.

## Contribution statement

AE conceptualized the design, AE and ATG generated and analyzed the data, AE, ATF, EBJ and JP interpreted the data and wrote the article.

## Data availability

All codes, structure files, and input parameters are available online under at https://github.com/agattar/Attar_ElasticShellModel.git. For further access to data relevant to this report,directly contact the corresponding authors.

