## Supplementary text for "Chromatin phase separation and nuclear shape fluctuations are correlated in a polymer model of the nucleus"

**The Molecular Dynamics Simulation Parameters**

The Lennard-Jones (LJ) potential was employed for modeling the non-bonded pairwise interaction between the beads. LJ potential was shifted and truncated as in the following,

$$V(r) = \{u\left[\left(\frac{\sigma}{r}\right)^9 - \left(\frac{\sigma}{r}\right)^6 + v_s\right] \quad r \leq r_c \quad 0 \quad\quad r > r_c$$

The cutoff distance was set as $r_c = 2.5\sigma$ where the σ is the unit length in our model, and the shift factor in the potential as $v_s = 0$. The attraction strengths used in the simulations are defined in **Table 1**. The interactions in the model are repulsive interactions. The strength of the repulsive interactions for the beads was set to $u = 1kT$ with a cutoff distance $r_c = 2^{\frac{1}{6}}\sigma$ and a shift factor to $v_s = \frac{1}{4}$. Unless otherwise noted, the timestep in the simulations was $\Delta t = 0.005\tau$, where τ is the unit of time. We employed the Langevin thermostat in our simulations with the microcanonical ensemble (*NVE)* to implement Brownian dynamics. The temperature has remained constant, and the damping coefficient was set to 1000τ unless stated otherwise. The periodic boundaries were used in all directions, and the dimensions of the simulation box were set as $60\times60\times60\sigma^3$. All simulations are run using LAMMPS MD package.

For the bonded interactions, we employed two approaches. Except for the bonds used for crosslinking between constitutive heterochromatin beads, all other assigned bonds followed the non-extensible FENE potential.

$$V(r) = -0.5kR_0^2 ln[1 - \left(\frac{r}{R_0}\right)^2]$$

The maximum bond distance for the potential was defined as $R_0$ =1.5, which defines the maximum extension that the FENE bond could have. The bond energy was set to $k = 5\frac{kT}{\sigma^2}$ for the shell beads and the bond energy between chromatin polymer was $k = 30\frac{kT}{\sigma^2}$. For the crosslinking of the constitutive heterochromatin, we used the harmonic bonds described,

$$V(r) = k(r - r_0)^2$$

where $r_0$= 1.5σ and the energy of the bond was set to $k = 10\frac{kT}{\sigma^2}$.

The flexibility of the polymer was provided using the harmonic angle potential. Heterochromatin fibers possess longer persistence lengths than euchromatin, within the range of 100-200 nm and 10-50 nm, respectively. Hence, the angles were only defined for the heterochromatin beads in our model, and the potential employed is the following,

$$V(\theta) = k_\theta(\theta - \theta_0)^2$$

where the energy, $k_\theta$, was set to $k_\theta = 1\frac{kT}{rad^2}$ and the angle was defined as $\theta = 180$ to obtain a semiflexible heterochromatin.

The initial shell size was determined to be 42σ. During the minimization simulation, the shell was shrunk to approximately 90% of the initial radius, resulting in a final radius of ~38σ. Furthermore, in the equilibrium simulations where the polymers were allowed to relax, the shell radius was further reduced to about 70% of the initial radius, resulting in a final radius of ~32σ. Finally, in the conventional simulations, the shell radius was ~28σ, which represents approximately 65% of the initially determined radius.

**The Details on Analysis of the Data**

*The Determination of the Chromatin Density*

The three various chromatin densities were employed in our simulations, and to do that, the number of triblock copolymers was decreased from $N_{polymer} = 8$ to $N_{polymer} = 6$ and $N_{polymer} = 4$. The final volume fractions were obtained from the end-product of the default conventional nucleus simulations. The following equation was utilized for calculating the chromatin density. $V_{shell} = \frac{4}{3} \times \pi \times < R_{sphere}>^3$

$$V_{chromatin} = N_{chromatin} \times (\frac{4}{3} \times \pi \times R_{chromatin}^3)$$

$$\varphi = \frac{V_{chromatin}}{V_{shell}}$$

where,

$$N_{chromatin} = N_{polymer} \times 6002$$

$$R_{chromatin} = 0.5\sigma$$

$R_{sphere}$ denotes for the average radial distance of the shell where $R_{chromatin}$ is the radius of the each bead in the model.

*The Measurement of Radial Distance and Density Profiles*

The radial distance of the chromatin beads was calculated by measuring the Euclidean distance from the center of the sphere at each simulation step, $t = 4\times 10^3$, and taking the average across the same bead types. The density profile of the chromatin beads was generated step-by-step with a constant increment rate as in the following.

$$V = \frac{4}{3} \times \pi \times \Delta R^3$$

The beads occupied in the defined volume were considered accordingly in the volume range by the following equation.

$$\Delta R - \Delta R_{step} < r_{bead} < \Delta R$$

Thus, the density profiles were generated in the defined volume by dividing the number of beads with the same type, which occupied the specified volume, by the total number of same-type beads.

$$\rho = \frac{N_{bead}}{N_{total}}$$

*The Analysis of the Nuclear Shape Fluctuations of the Shell*

The shape's fluctuations were calculated by taking a thin slice, having a width of $w$=1σ from the x-axis in our model. Since our shell rotates around its center, a different number of beads are obtained at each slice. The Fast Fourier Transform (FFT) algorithm was run on the 230 modes, q; hence, the 230 beads on that slice were randomly selected and ordered by their angle from the center. The algorithm was run using the radial distances of those beads chosen from the center, and the fluctuations were calculated at each step after the phase separation was completed, which is the last half period of the simulations. As an additional measure of bead fluctuations, the root-mean-squared distance of each bead was calculated. The standard deviation from the radial distances was averaged at each time step during the last half period of the simulations.

| Type | Interaction Energy |
|---|---|
| Euchromatin - Euchromatin | 0.05 kT |
| Euchromatin - Facultative Heterochromatin | 0.25 kT |
| Euchromatin – Cons. Heterochromatin | 0.50 kT |
| Fac. Heterochromatin – Fac. Heterochromatin | 0.65 kT |
| Heterochromatin-Shell | 0.75 kT |
| Fac. Heterochromatin – Cons. Heterochromatin | 0.85 kT |
| Cons. Heterochromatin – Cons. Heterochromatin | 1.10 kT |
| Telomere-Shell | 2.20 kT |

**Table 1** The interaction energies set for various beads in the simulations where k and T are Boltzmann and absolute temperature.

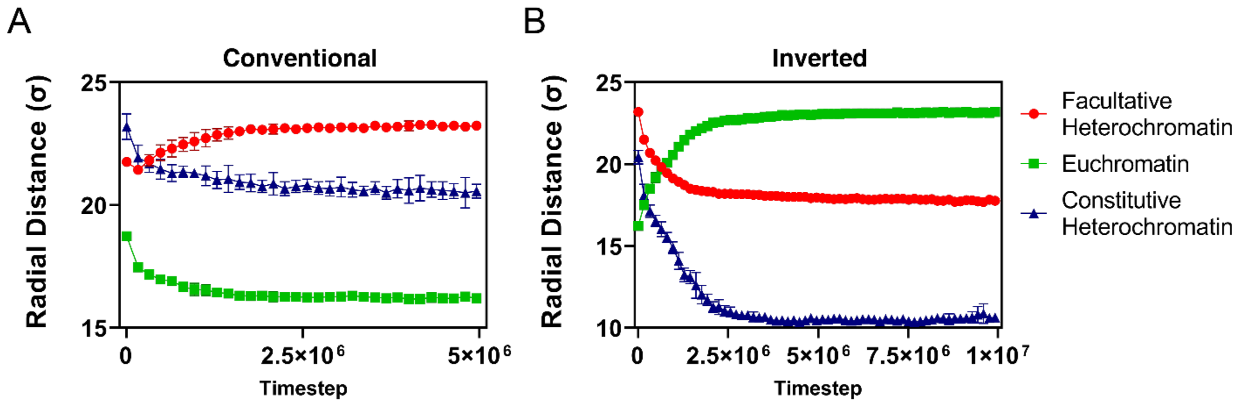


FIG. S1. Each chromatin domain's time-dependent radial distribution profiles in conventional and inverted nucleus simulations. The radial distance of the each bead that has the same type is calculated and the mean radial distance is measured.

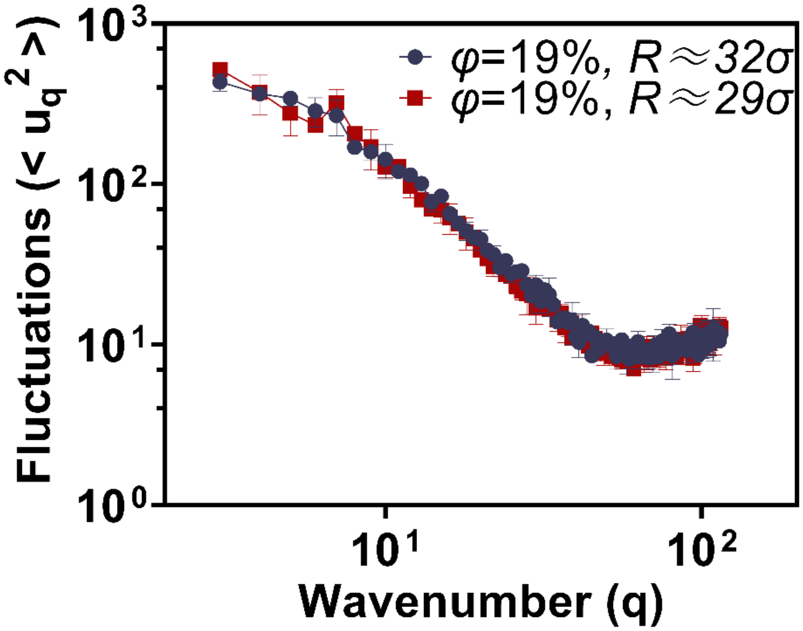


FIG. S2. The nuclear shape fluctuations of the two models have the same chromatin density with different radial sizes of model nucleus. Each case has different number of shell beads however have the same number of polymer beads. For the case where $R$=32σ, number of shell beads is $N$=18000 and for the other case, $N$=15000.

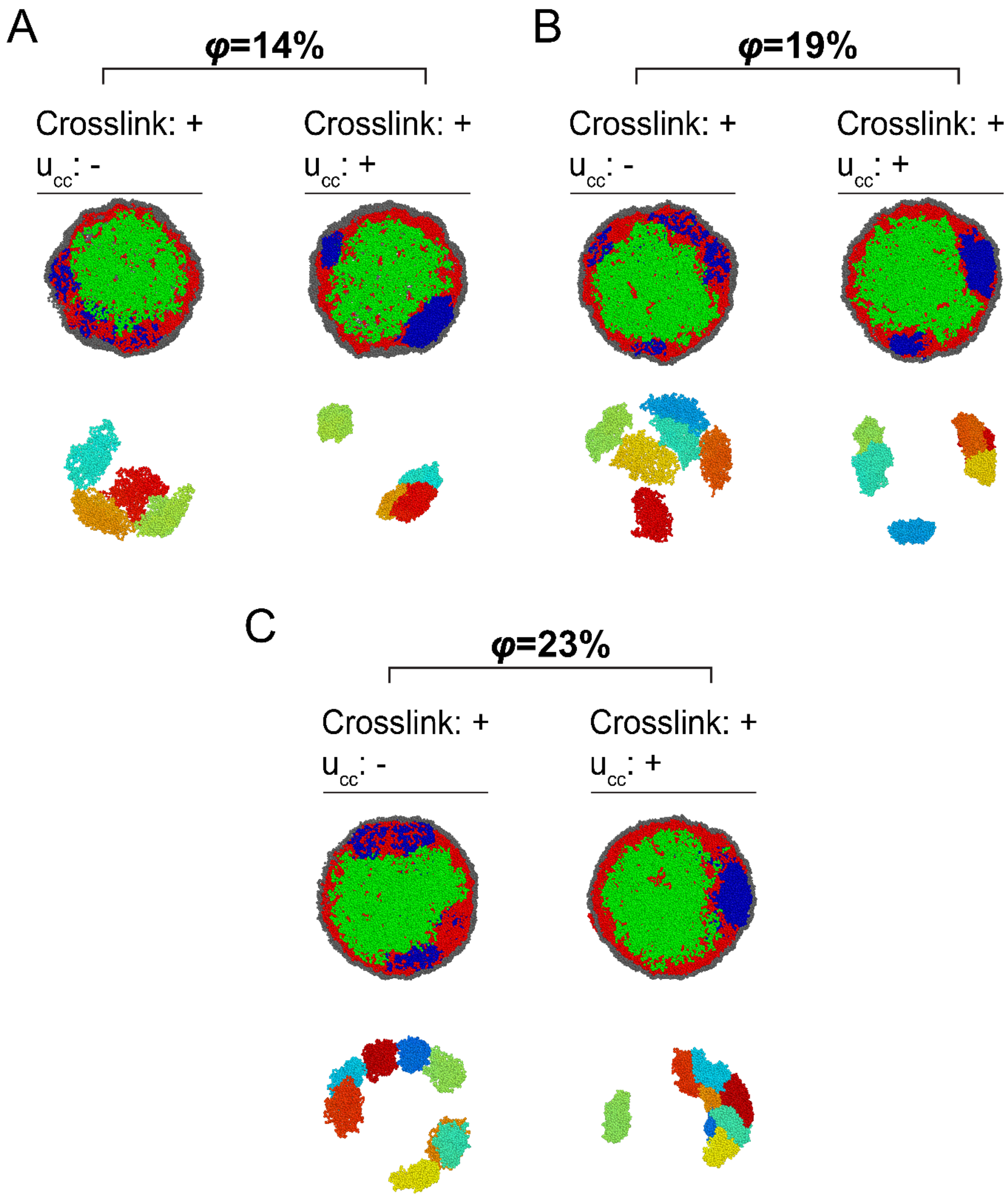


FIG. S3. The representative snapshots are taken from the simulation at the last time step for each triblock polymer under three different volume fractions, respectively. The attractions between constitutive heterochromatin were either turned on or off for each case and crosslinks are implemented via harmonic bonds. Lower panels in A-C, shows the constitutive heterochromatin domain only. Constitutive heterochromatin domain of each chromosome donated by a different color in the snapshots.

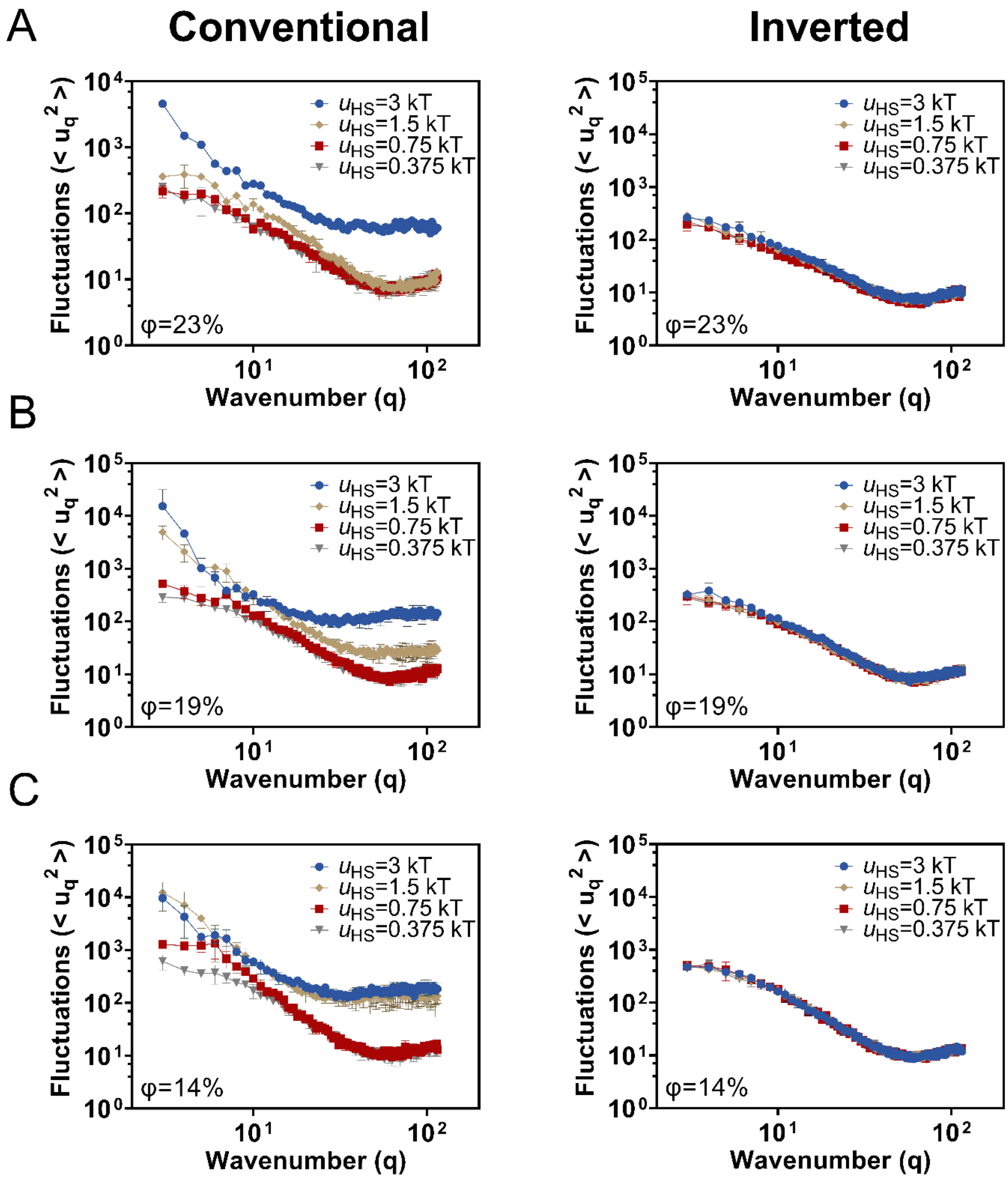

FIG. S4. The nuclear shape fluctuations extracted with various chromatin densities from conventional and inverted nucleus simulations **A)** $\varphi$=23%, **B)** $\varphi$=19%, and **C)** $\varphi$=14%. The interactions of facultative and constitutive heterochromatin with the shell are only altered in these cases.

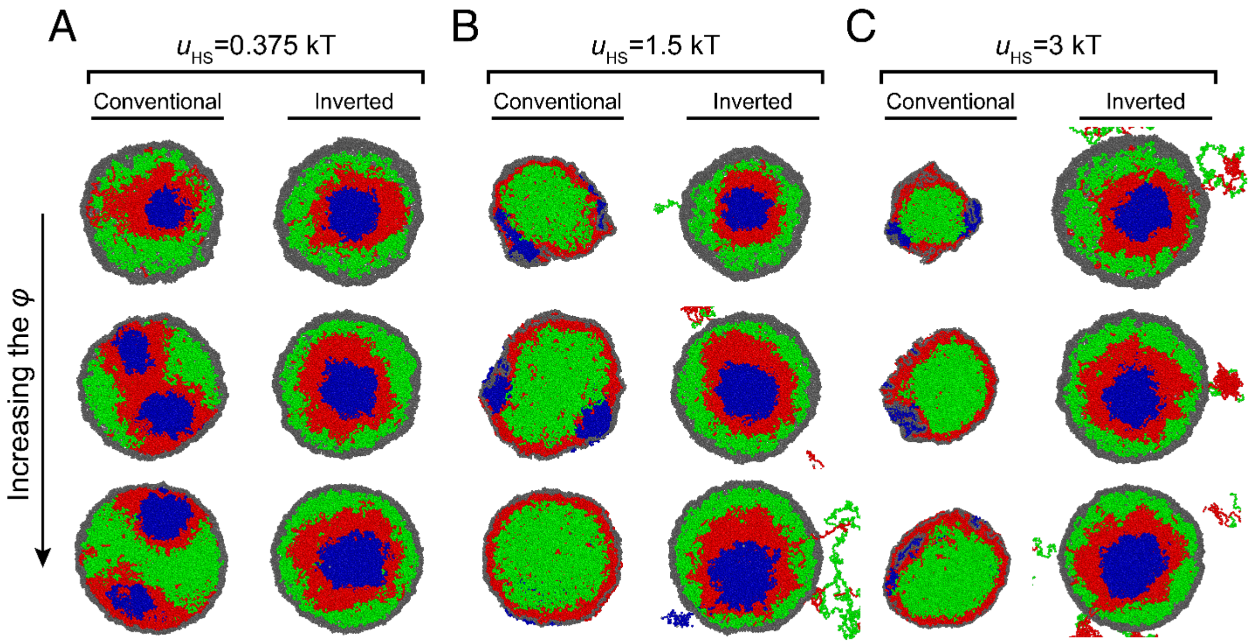


FIG. S5. The snapshots from our model under different chromatin densities and heterochromatin-shell, $u_{HS}$, interactions. The green, blue and red beads correspond to euchromatin, constitutive heterochromatin and facultative heterochromatin, respectively. For the inverted nucleus, all the chromatin-shell interactions are depleted.

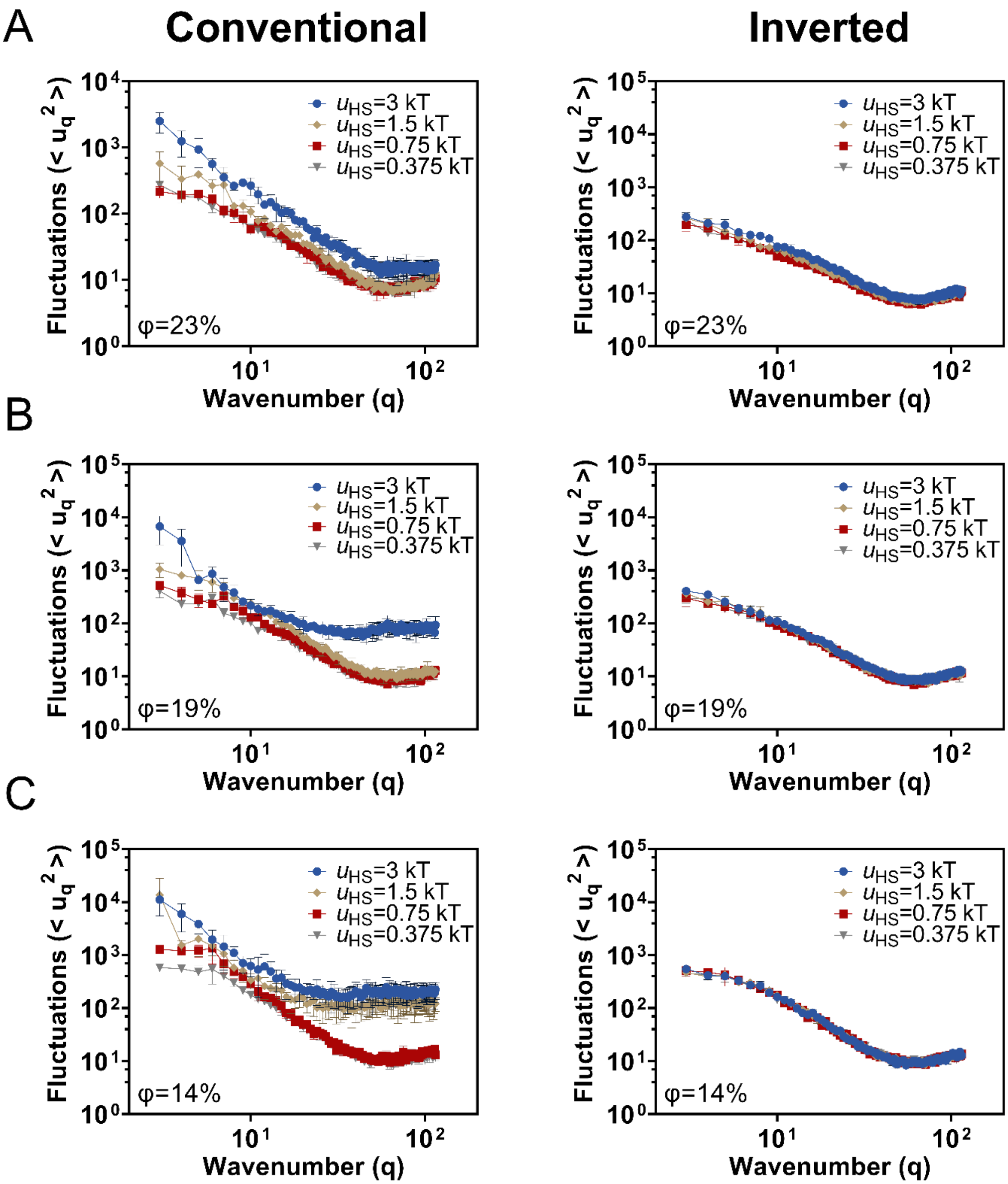


FIG. S6. The nuclear shape fluctuations extracted with various chromatin densities from conventional and inverted nucleus simulations **A)** $\varphi$=23%, **B)** $\varphi$=19%, and **C)** $\varphi$=14%. The interactions between facultative heterochromatin and the shell are only altered in these cases.

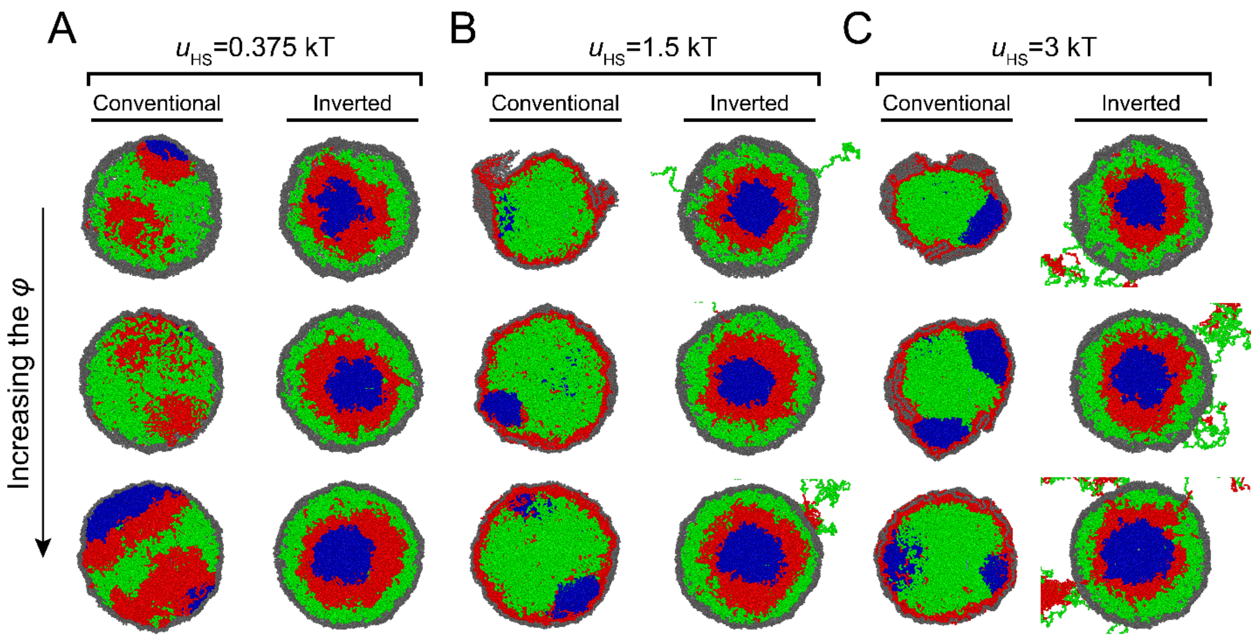


FIG. S7. The snapshots from our model under different chromatin densities and facultative heterochromatin-shell, $u_{\mathrm{HS}}$, interactions. The green, blue and red beads correspond to euchromatin, constitutive heterochromatin and facultative heterochromatin, respectively. For the inverted nucleus, all the chromatin-shell interactions are depleted.

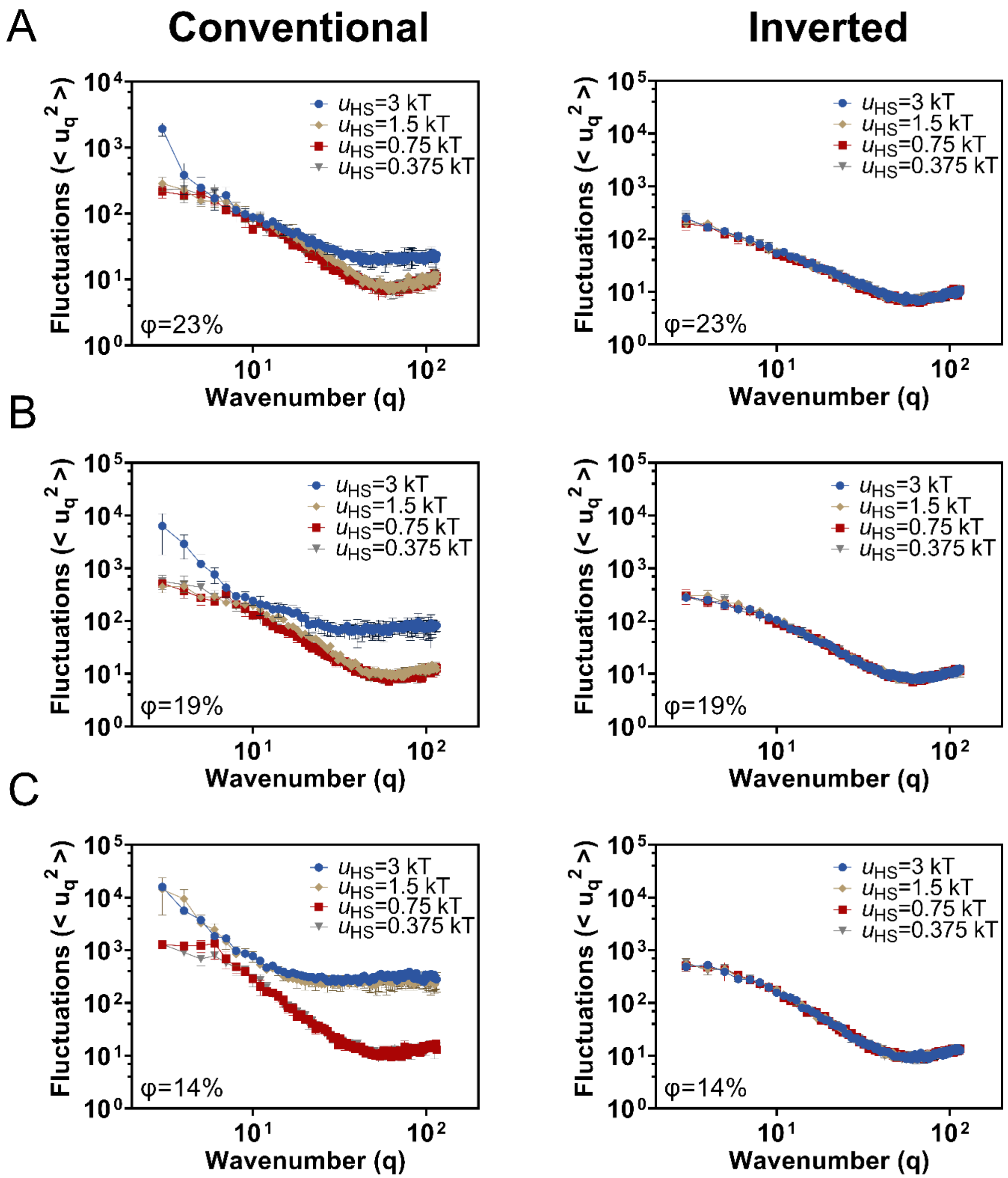


FIG. S8. The nuclear shape fluctuations extracted with various chromatin densities from conventional and inverted nucleus simulations **A)** $\varphi$=23%, **B)** $\varphi$=19%, and **C)** $\varphi$=14%. The interactions between constitutive heterochromatin and the shell are only altered in these cases.

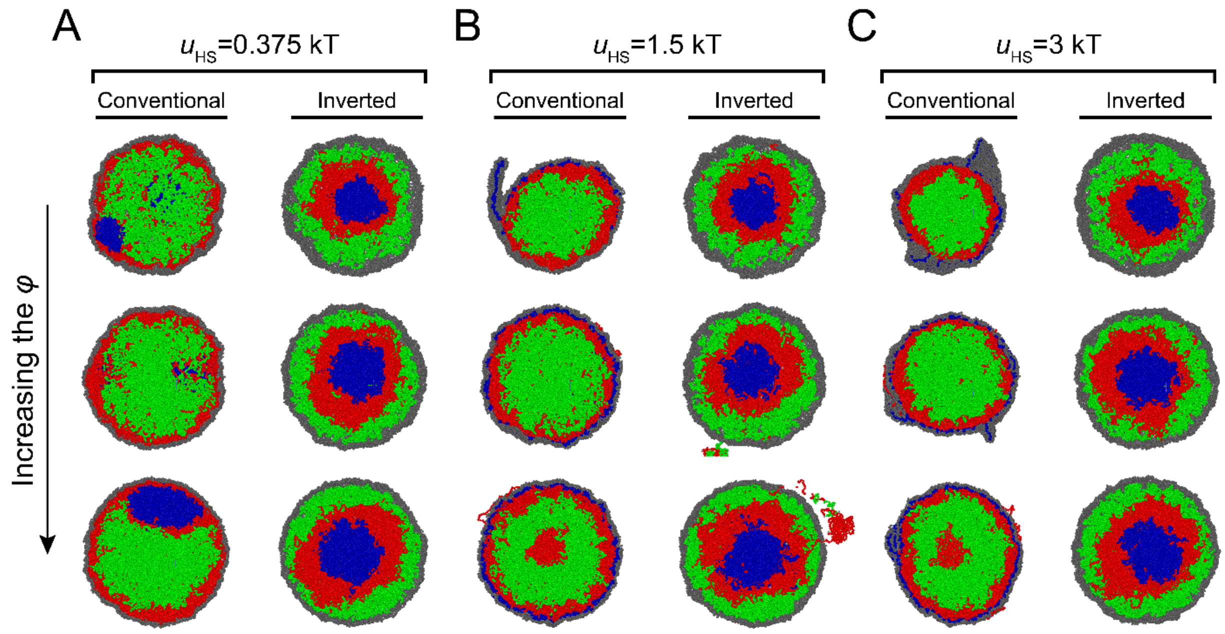


FIG. S9. The snapshots from our model under different chromatin densities and constitutive heterochromatin-shell, $u_{HS}$, interactions. The green, blue and red beads correspond to euchromatin, constitutive heterochromatin and facultative heterochromatin, respectively. For the inverted nucleus, all the chromatin-shell interactions are depleted.

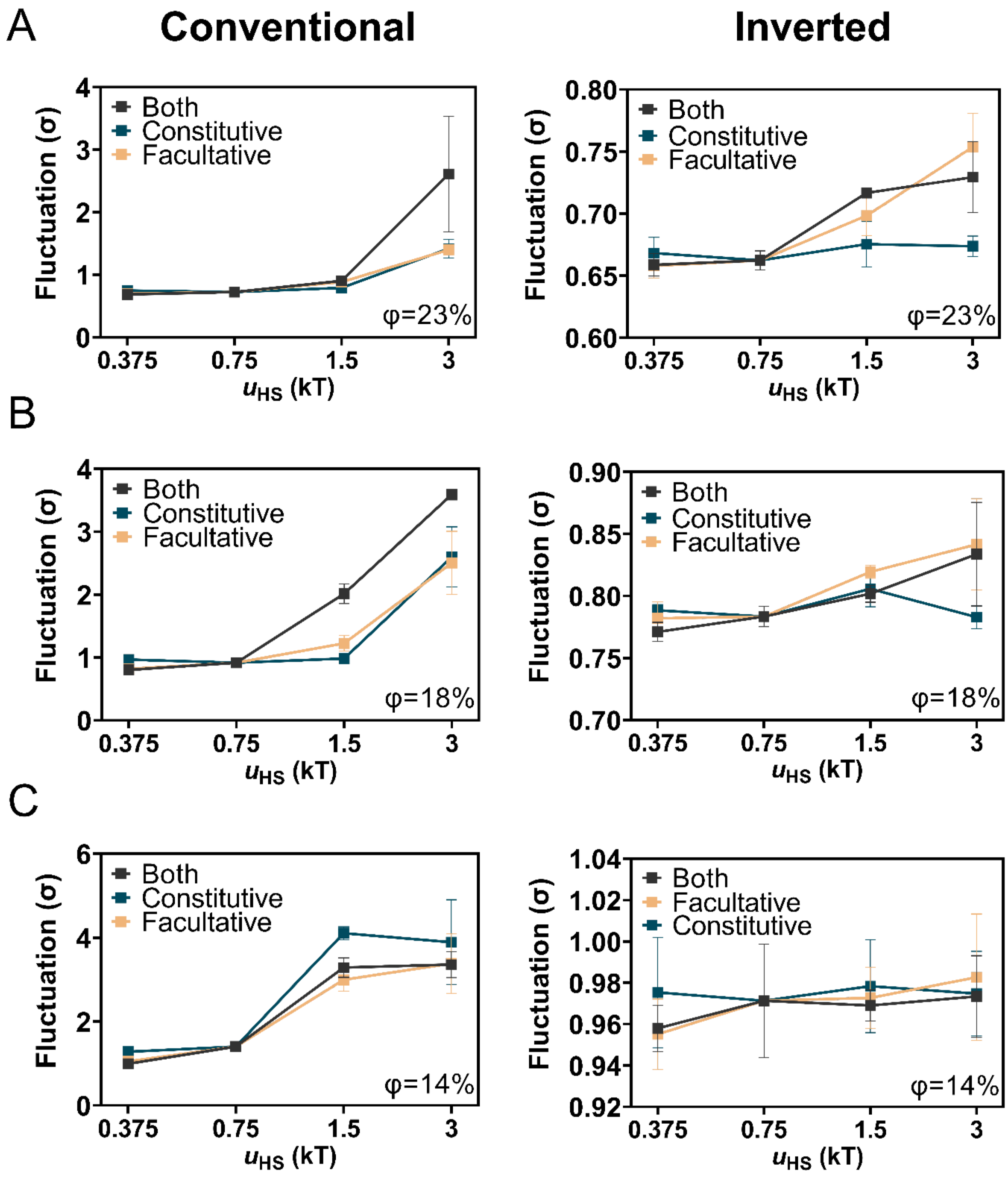


FIG. S10. The RMS radial fluctuations of different heterochromatin-shell interaction strengths of conventional and inverted nucleus simulations under three volume fractions: **A)** $\varphi$=23%, **B)** $\varphi$=19%, and **C)** $\varphi$=14%.

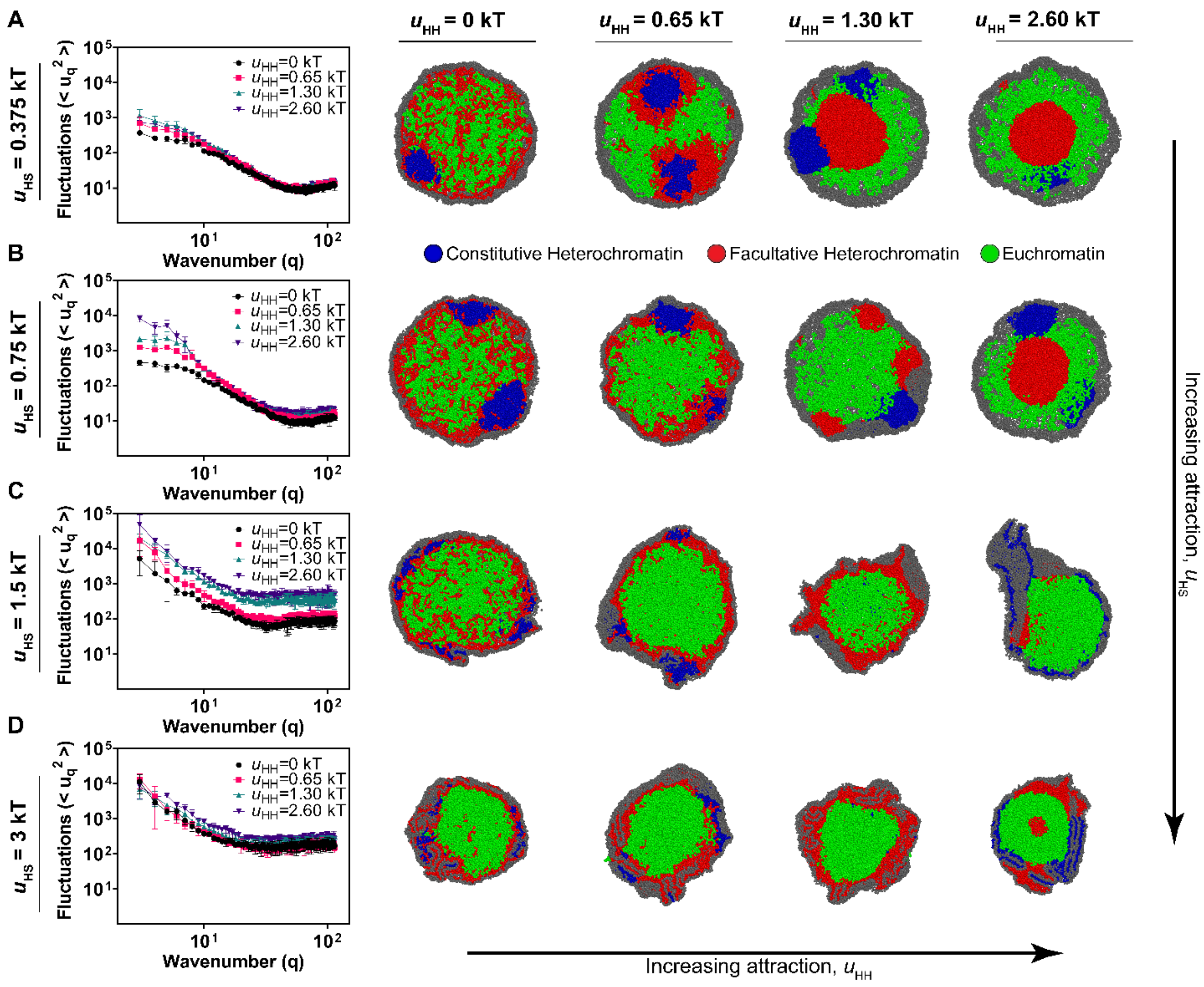


FIG. S11. The nuclear shape fluctuations and the snapshots from the conventional nucleus simulations with various heterochromatin-shell interactions and self-facultative heterochromatin attractions **A)** $u_{HS}$=0.375 kT, **B)** $u_{HS}$=0.75 kT, **C)** $u_{HS}$=1.50 kT and **D)** $u_{HS}$=3 kT. The chromatin density was set to $\varphi$=14%.

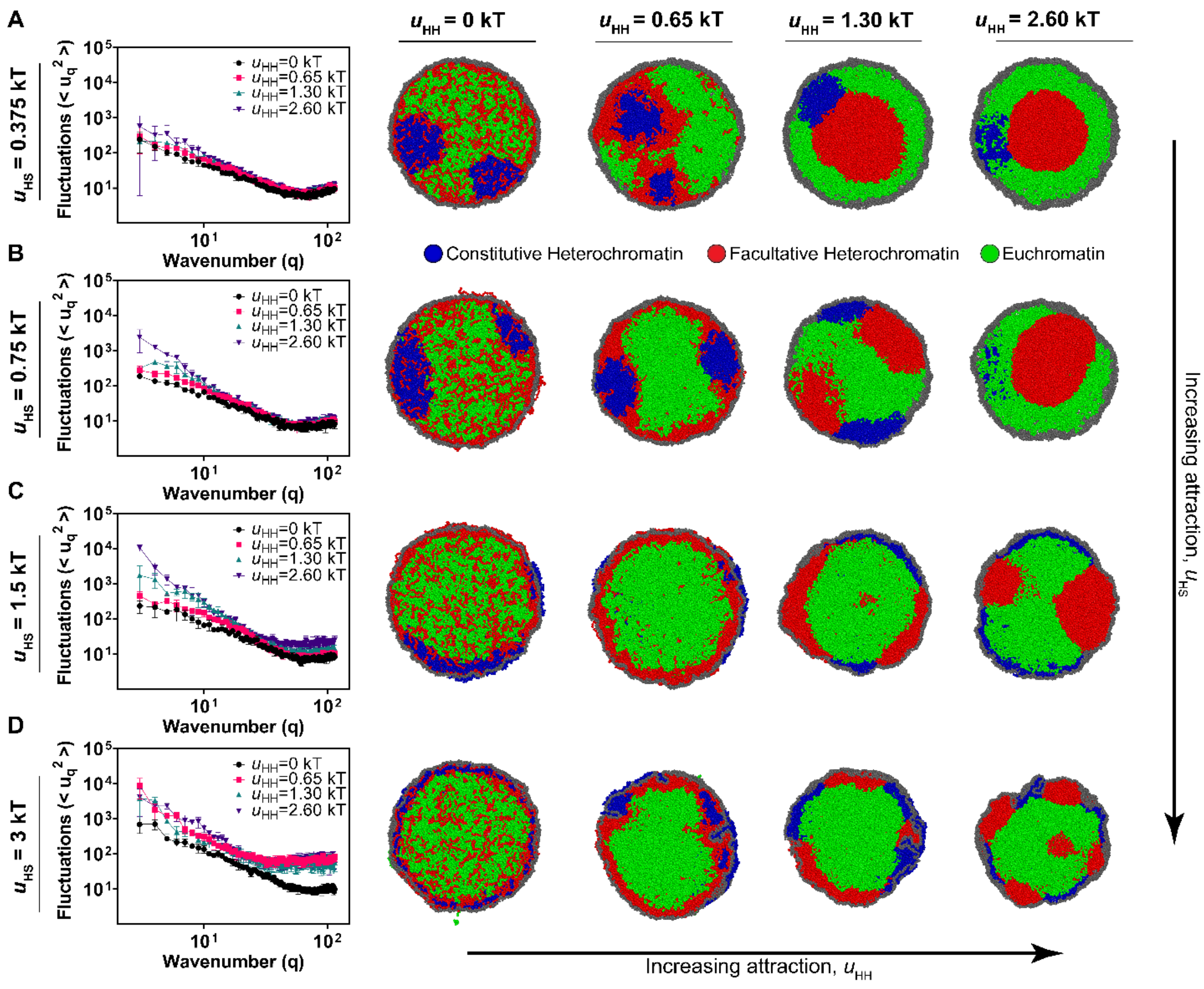


FIG. S12. The nuclear shape fluctuations and the snapshots from the conventional nucleus simulations with various heterochromatin-shell interactions and self-facultative heterochromatin attractions **A)** $u_{HS}$=0.375 kT, **B)** $u_{HS}$=0.75 kT, **C)** $u_{HS}$=1.50 kT and **D)** $u_{HS}$=3 kT. The chromatin density was set to $\varphi$=23%.

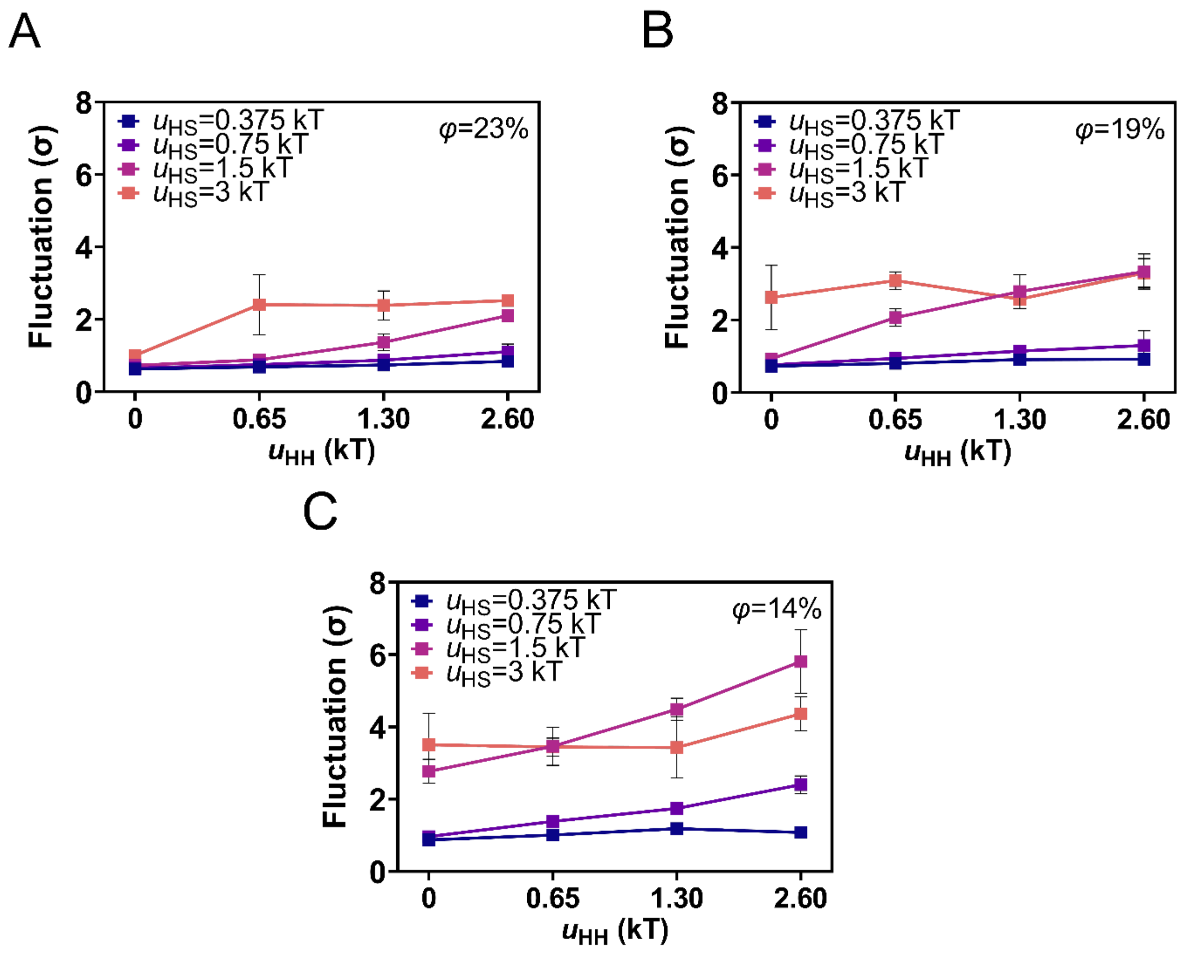


FIG. S13. The RMS radial fluctuations for the conventional nucleus simulations under three different chromatin densities, **A)** $\varphi$=23%, **B)** $\varphi$=19%, and **C)** $\varphi$=14%.

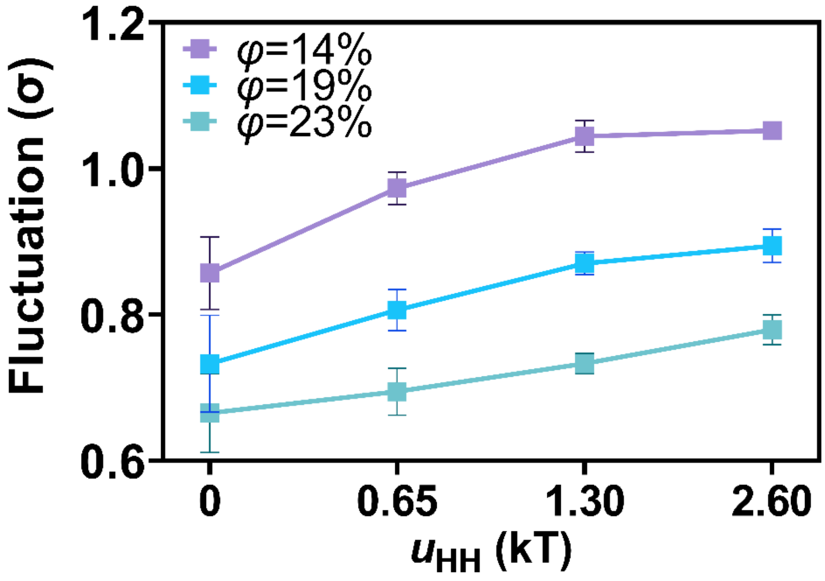


FIG. S14. The RMS radial fluctuations of inverted nucleus as a function of various facultative heterochromatin, $u_{HH,}$, self-attractions under three different volume fractions. Note that, in inverted nucleus heterochromatin-shell interactions are depleted.

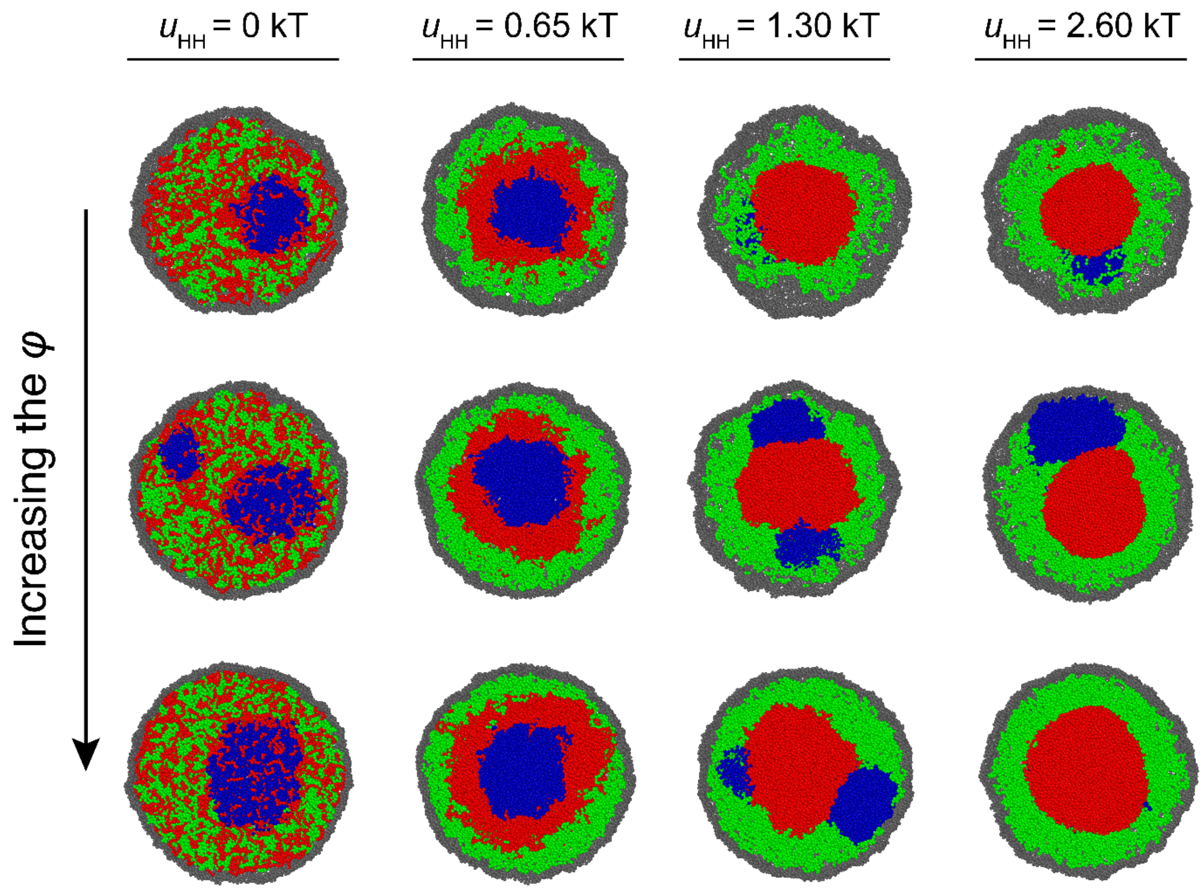


FIG. S15. The snapshots from our inverted nucleus model under different chromatin densities and facultative heterochromatin self-attraction, $u_{\mathrm{HH}}$. The green, blue and red beads correspond to euchromatin, constitutive heterochromatin and facultative heterochromatin, respectively. For the inverted nucleus, all the chromatin-shell interactions are depleted.

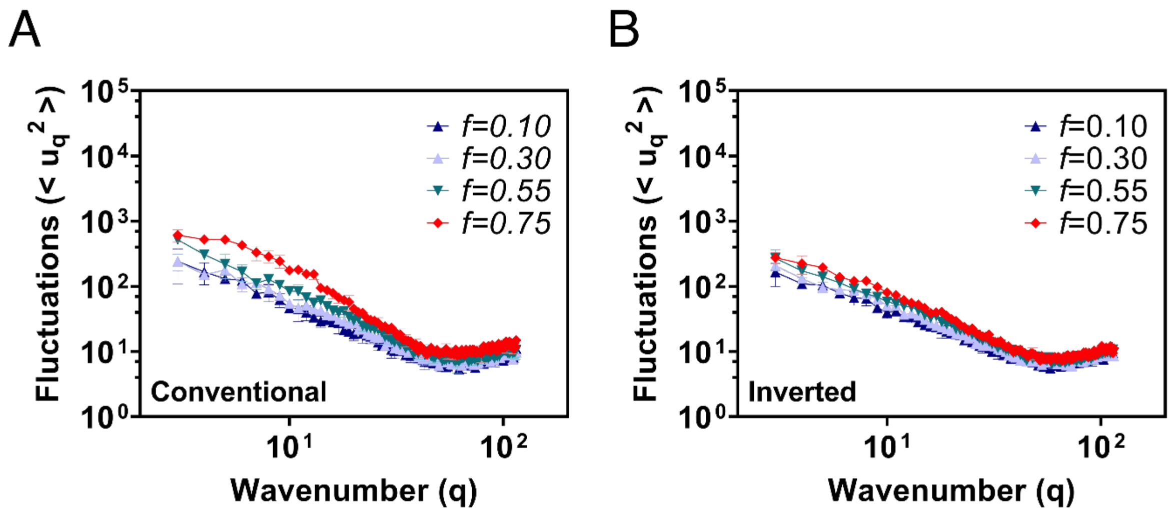


FIG. S16. The shape's fluctuations for various facultative heterochromatin fractions under chromatin density of $\varphi$=23% from **A)** conventional nucleus and **B)** inverted nucleus simulations.

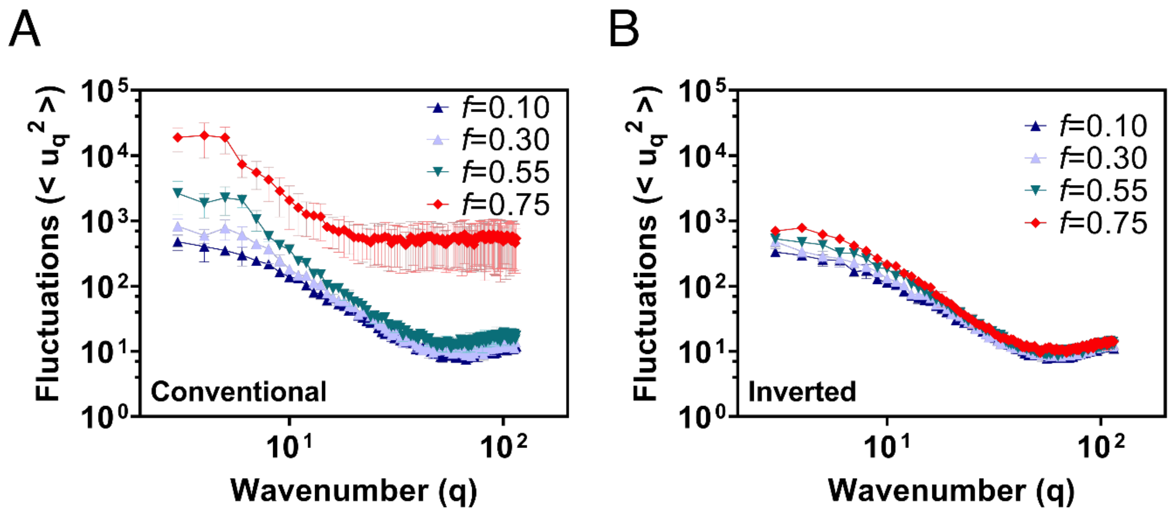


FIG. S17. The shape's fluctuations for various facultative heterochromatin fractions under chromatin density of $\varphi$=14% from **A)** conventional nucleus and **B)** inverted nucleus simulations.

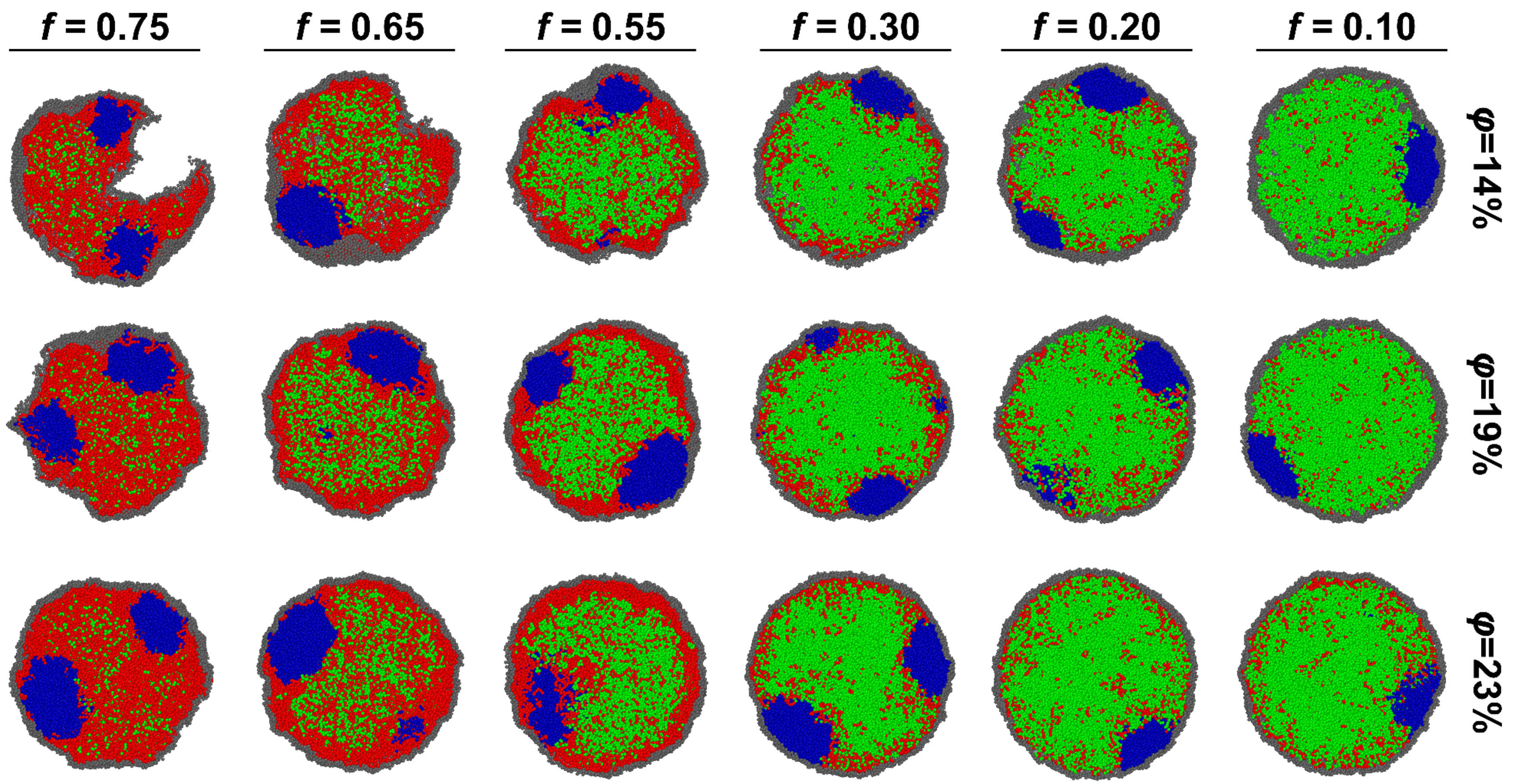


FIG S18 The snapshots from conventional nucleus organizations for various heterochromatin and volume fraction. The green, blue and red beads correspond to euchromatin, constitutive heterochromatin and facultative heterochromatin, respectively.

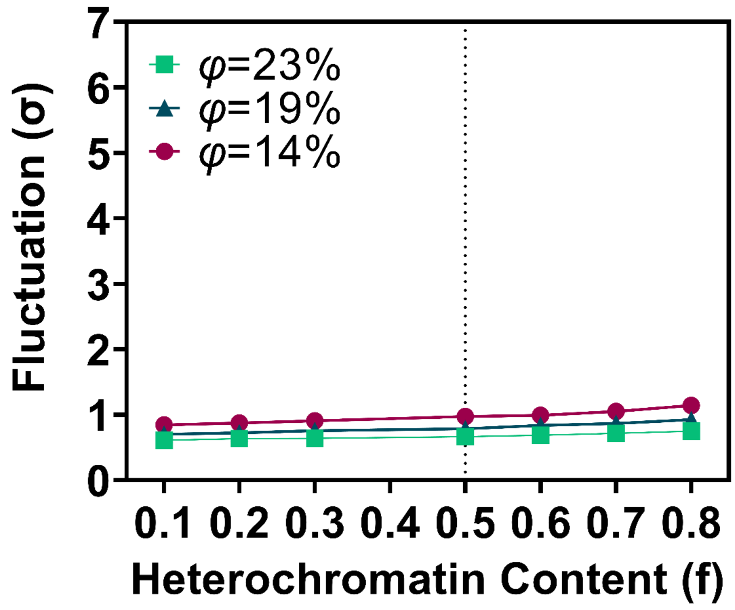


FIG. S19. The RMS radial fluctuations of inverted nucleus model as a function of heterochromatin fractions for three defined chromatin densities. Note that in inverted nucleus, heterochromatin-shell interactions, $u_{HS,}$ are depleted.

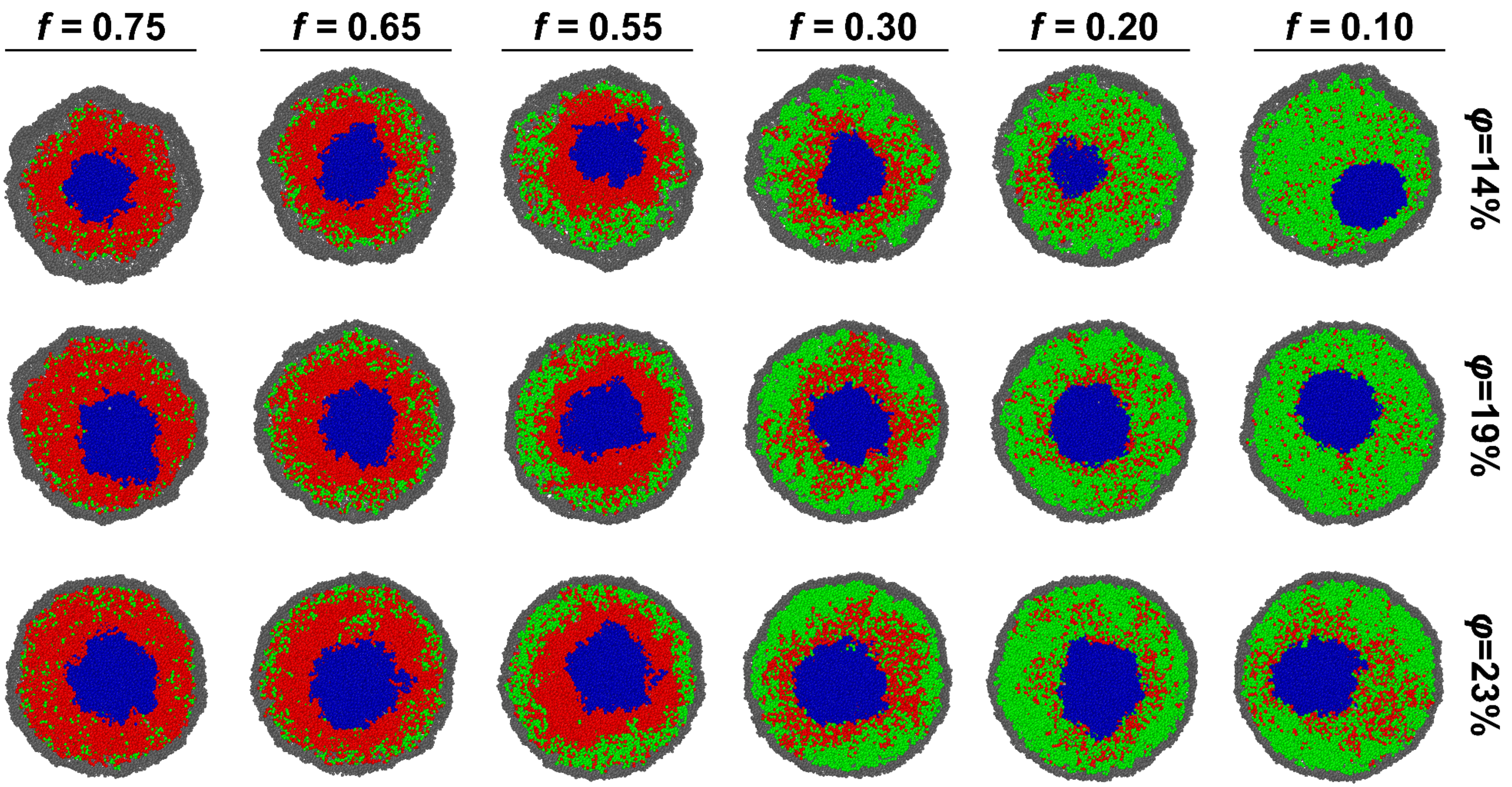


FIG S20 The snapshots from inverted nucleus organizations for various heterochromatin and for volume fractions. The snapshots from conventional nucleus organizations for various heterochromatin and volume fraction. The green, blue and red beads correspond to euchromatin, constitutive heterochromatin and facultative heterochromatin, respectively.

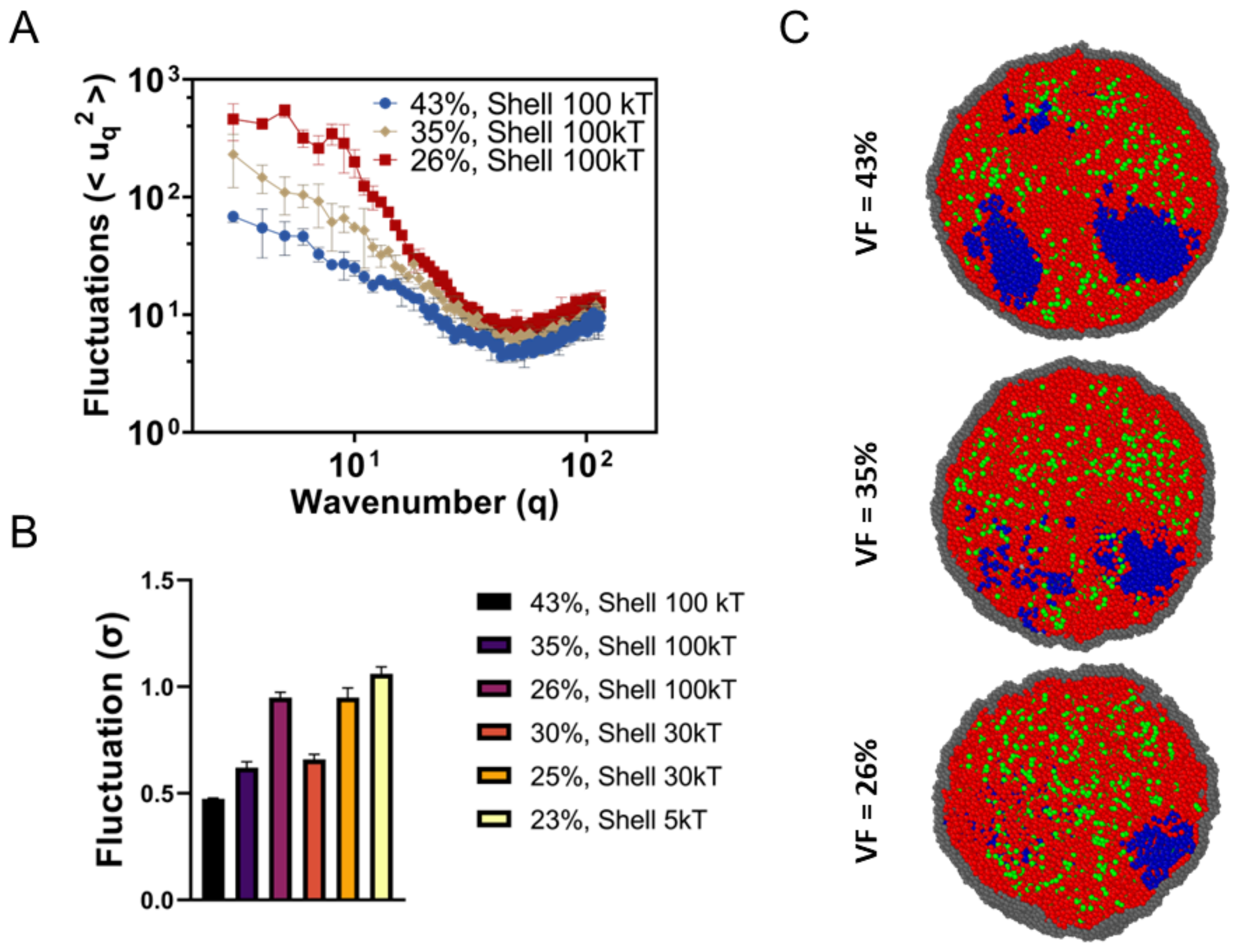


FIG. S21 The impact of the bond stiffness of the nuclear shape fluctuations. The heterochromatin fraction in this cases are set to $f$=0.75. **A)** The nuclear shape fluctuation of the various volume fractions having the same shell bond stiffness. **B)** The RMS radial fluctuations for different volume fractions and shell bond stiffness. **C)** The snapshots from the conventional simulation with various volume fractions. The green, blue and red beads correspond to euchromatin, constitutive heterochromatin and facultative heterochromatin, respectively.
